# The Cellular and Synaptic Actions of Dopamine During Behavior

**DOI:** 10.64898/2026.08.05.742858

**Authors:** Mélanie Druart, Yun C. Yang, Nicolas X. Tritsch, Tanya Sippy

## Abstract

Dopamine signaling in the striatum is essential for a wide range of functions, from reward learning to motor vigor and behavioral flexibility. Although dopamine signals fluctuate on sub-second timescales, how these signals are translated into lasting changes in striatal circuit function remains unknown. Resolving this requires cell-type-specific measurements of synaptic and intrinsic properties during behavior, a longstanding technical challenge. Here, we combined *in vivo* whole-cell membrane potential recordings, simultaneous monitoring and bidirectional manipulation of dopamine signaling in awake, behaving mice to examine how dopamine shapes corticostriatal circuits. Acute manipulations of dopamine over seconds to minutes produced only modest effects on corticostriatal synaptic transmission and no detectable changes in membrane potential dynamics or intrinsic excitability. By contrast, associative learning robustly strengthened identified corticostriatal synapses onto both D1- and D2-expressing spiny projection neurons, yet only D1-SPN plasticity required dopamine signaling. These findings challenge models in which dopamine acts rapidly to tune striatal excitability and identify learning related plasticity as its principal mechanism for shaping striatal circuits *in vivo*.

## Introduction

Dopamine transmission in the striatum is essential for behaviors spanning multiple timescales, from initiating and invigorating ongoing actions to shaping future behavior through reinforcement learning (*1*). Its loss or dysregulation underlies disorders as diverse as Parkinson’s disease, addiction, and schizophrenia (*2–4*). Yet despite decades of work, a fundamental question remains: through which physiological mechanisms does dopamine reshape striatal circuits to influence behavior (*5*)?

Dopamine has long been proposed to reinforce actions by modifying corticostriatal circuits (*6*). Seminal work demonstrated that pairing dopaminergic signals with cortical activity induces synaptic plasticity *in vivo*, establishing dopamine-dependent plasticity as a candidate cellular mechanism for reinforcement learning (*7*, *8*). Subsequent studies in acute brain slices defined the underlying physiological mechanisms, showing that dopamine receptor activation regulates both the intrinsic excitability of striatal spiny projection neurons and corticostriatal synaptic plasticity (*9–16*). Together, these studies established the prevailing physiological model in which dopamine shapes corticostriatal circuits through coordinated regulation of intrinsic excitability and synaptic plasticity.

Whether these mechanisms account for dopamine’s principal physiological actions during natural behavior, however, remains unresolved. In the intact brain, endogenous dopamine release fluctuates continuously with rewards, predictive cues, movement, and internal state rather than occurring as isolated experimental manipulations (*17–21*). Recent imaging studies further demonstrated that physiological dopamine transients rapidly engage intracellular PKA signaling that persists over behaviorally relevant timescales (*22*), yet how this signaling is translated into changes in neuronal physiology and synaptic plasticity remains unknown. At the same time, recent studies have questioned whether physiological dopamine transients produce the rapid modulation of neural activity or behavior predicted by prevailing models of acute intrinsic modulation (*23–25*), while revealing highly localized spatiotemporal dopamine signaling within striatal circuits (*26*). Consequently, it remains unclear through which physiological mechanisms, and over what timescales, endogenous dopamine shapes corticostriatal circuits during behavior.

To address this question, we combined extensive *in vivo* whole-cell membrane potential (V_m_) recordings (>300 neurons), cell-type-specific circuit identification, longitudinal learning paradigms, and simultaneous monitoring and bidirectional manipulation of dopamine signaling in awake, behaving mice. This integrated approach allowed us to directly determine how endogenous dopamine shapes neuronal physiology and synaptic plasticity during behavior.

## Results

### An *in vivo* approach for measuring dopamine actions at identified corticostriatal synapses

To directly determine how dopamine influences corticostriatal synapses *in vivo*, we developed an experimental preparation that allowed independent manipulation of cortical input and dopamine release while performing whole-cell recordings from genetically identified striatal projection neurons (Fig. 1A). We expressed the red-shifted opsin Chrimson in layer 5 neurons of primary somatosensory cortex (S1) and performed membrane potential (V_m_) recordings from neurons in the dorsolateral striatum (DLS) of awake, head-fixed D2-Cre × Ai32 × DAT-Flp mice, which express ChR2 in D2-SPNs and Flp recombinase in dopaminergic neurons (Fig. 1, B to C). D2-SPNs were identified by optogenetic tagging (optopatch), while unlabeled neurons were classified as putative D1-SPNs based on the absence of light-evoked responses (Fig. S1A; Table S2). SPNs had intrinsic properties typical of these neurons *in vivo* (*27*, *28*), with D2-SPNs showing greater excitability at baseline (Fig. S1B). Corticostriatal connections were identified as short-latency EPSPs evoked by optical stimulation of S1 axons and was observed in both D1- and D2-SPNs, permitting direct physiological measurements from identified corticostriatal synapses *in vivo* (Fig. 1, C to D).

**Fig. 1.**
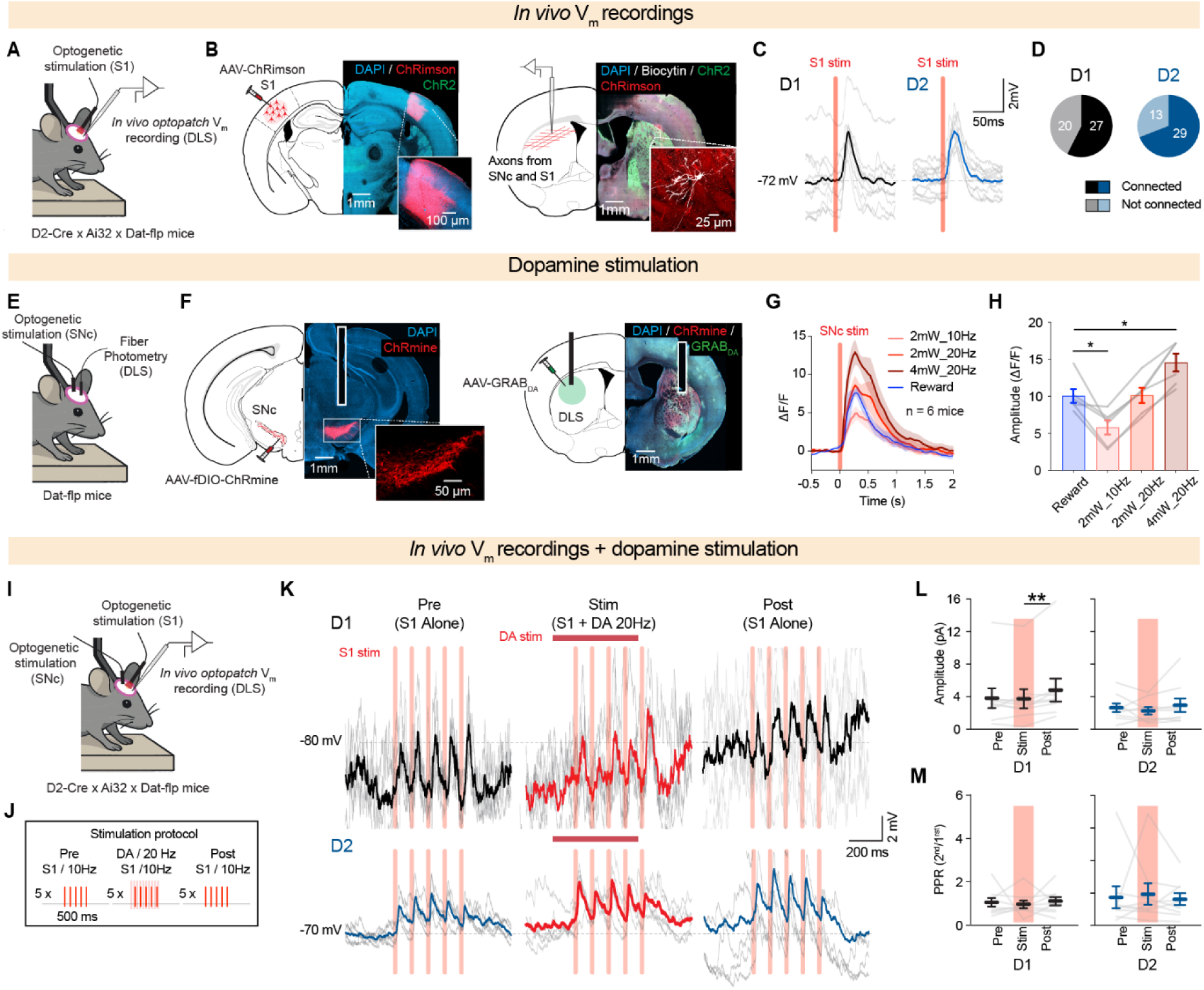
An *in vivo* membrane potential recording platform to investigate the physiological actions of dopamine. **(A)** Schematic of the recording configuration. AAV-ChRimson was expressed in S1 of D2-Cre × Ai32 × DAT-Flp mice, allowing optical control of cortical inputs through the skull. Whole-cell membrane potential (V_m_) recordings were performed allowing access to the membrane potential (V_m_) of neurons in the DLS of head-fixed awake mice. **(B)** Left: representative histology showing viral expression of AAV-ChRimson in S1 cortex (scale bars 1 mm and 50 μm). Left: a biocytin-filled neuron recorded in the DLS showing co-expression of ChR2 in the striatum together with ChRimson axons from S1. **(C)** Representative traces of S1 stimulation in D1-SPNs (top) and D2-SPNs (bottom) at 10 Hz, 4 mW. **(D)** Proportion of D1-SPNs (left) and D2-SPNs (right) receiving S1 input, as confirmed by short-latency EPSPs following cortical stimulation (D1: n = 27 cells connected, n = 20 cells not connected; D2: n = 29 cells connected, n = 13 cells not connected). **(E)** Schematic of the fiber photometry configuration for dopamine stimulation validation. AAV-fDIO-ChRmine was expressed in SNc dopamine neurons of DAT-Flp mice and AAV-GRAB-DA was expressed in the DLS with a fiber optic cannula placed over the DLS. **(F)** Representative histology showing viral expression of AAV-fDIO-ChRmine in SNc dopamine neurons, and ChRmine and GRAB-DA co-expression in the DLS with fiber optic placement. **(G)** Average fiber photometry trace of dopamine release in the DLS aligned to natural reward delivery (blue) and during different light stimulation parameters (LED 635 nm, 500 ms pulse; 2 mW 10 Hz, 2 mW 20 Hz, 4 mW 20 Hz; n = 6 mice. **(H)** Peak dopamine amplitude across stimulation conditions compared to natural reward delivery (repeated measures One-way ANOVA, *P* = 0.0002; n = 6 mice), confirming that 2 mW at 20 Hz most closely approximates the reward-evoked dopamine transient. **(I)** Schematic of the recording configuration to assess S1 DLS synaptic strength modulated by dopamine stimulation. AAV-fDIO-ChRmine was expressed in SNc dopamine neurons and AAV-ChRimson was expressed in S1 of D2-Cre x Ai32 x DAT-Flp mice, allowing independent optical control of cortical inputs and dopamine release. V_m_ recordings were performed in the DLS of head-fixed awake mice. **(J)** Schematic of the stimulation protocol. Five repetitions of 5 S1 stimulations at 10 Hz were delivered with 10 seconds intervals before, during paired S1 and SNc dopamine neuron stimulation (20 Hz), and after SNc stimulation. **(K)** Representative traces of S1 stimulation in D1-SPNs (top) and D2-SPNs (bottom) before, during and after dopamine stimulation. Individual trials shown in gray, average in color. **(L)** EPSP amplitude before, during, and after paired with dopamine stimulation in D1-(left) and D2-SPNs (right, repeated measures One-way ANOVA, D1: *P* = 0.006, n = 9 cells from 6 mice; D2: *P* = 0.40 n = 9 cells from 6 mice). **(M)** Paired-pulse ratio of the second stimulation before, during, and after paired with dopamine stimulation in D1- and D2-SPNs (repeated measures One-way ANOVA, D1: *P* = 0.86, n = 9 cells from 6 mice; D2: *P* = 0.82, n = 9 cells from 6 mice). Data are shown as mean ± SEM. Each open circle represents an individual neuron. \**P* < 0.05, \*\**P* < 0.01.

To independently manipulate dopamine signaling, we expressed the excitatory opsin ChRmine selectively in substantia nigra pars compacta (SNc) dopamine neurons (Fig. 1E-F, fig. S1, C and D). Fiber photometry using the green dopamine sensor GRABDA2m confirmed that optical stimulation reliably evoked dopamine transients in the DLS, and stimulation parameters were calibrated to match the amplitude of dopamine release observed following natural reward delivery (Fig. 1G-H; Table S1). This preparation enabled independent control of corticostriatal input and dopamine release while monitoring postsynaptic responses from identified SPNs.

Having established independent control of S1 inputs and dopamine release together with cell-type-specific recordings from connected SPNs, we next combined these elements in a single *in vivo* preparation. In connected D1- and D2-SPNs, S1 stimulation was delivered before, during, and after concurrent activation of SNc dopamine neurons (Fig. 1, I and J). Acute dopamine release produced a small but significant increase in EPSP amplitude after the SNc stimulation in D1-SPNs, whereas D2-SPNs showed no detectable change (Fig. 1, K and L). Paired-pulse ratio was unchanged in either cell type (Fig. 1M), suggesting that the modest enhancement of synaptic transmission reflected a postsynaptic mechanism. Together, these experiments establish an approach for directly measuring dopamine actions at identified corticostriatal synapses *in vivo* and demonstrate that physiological dopamine exerts modest acute effects on corticostriatal synaptic transmission over short timescales.

### Acute dopamine does not rapidly alter intrinsic excitability *in vivo*

Dopamine has also been proposed to influence striatal function by altering the intrinsic excitability of SPNs. To directly test this possibility *in vivo*, we optogenetically activated SNc dopamine neurons while performing whole-cell recordings from optically identified D2-SPNs and putative D1-SPNs in awake D2-Cre × Ai32 × DAT-Flp mice (Fig. 2A). To assess intrinsic excitability, we injected depolarizing current ramps and measured rheobase, defined as the minimum current required to evoke an action potential (Fig. 2B). After establishing baseline rheobase, a 20-Hz, 500-ms light pulse was delivered immediately before each current ramp to evoke dopamine release during membrane depolarization (Fig. 2C). Control recordings were performed in mice lacking ChRmine expression to account for repeated depolarization alone. Acute dopamine release produced no detectable change in rheobase in either D1- or D2-SPNs compared with controls (Fig. 2, D and E; Table S3). Consistent with this, bidirectional manipulation of dopamine signaling produced no detectable changes in ongoing membrane potential dynamics, and spontaneous dopamine fluctuations were uncorrelated with membrane potential in either D1- or D2-SPNs when recorded simultaneously (Figs. S2 and S3; Table S4 and S5).

**Fig. 2.**
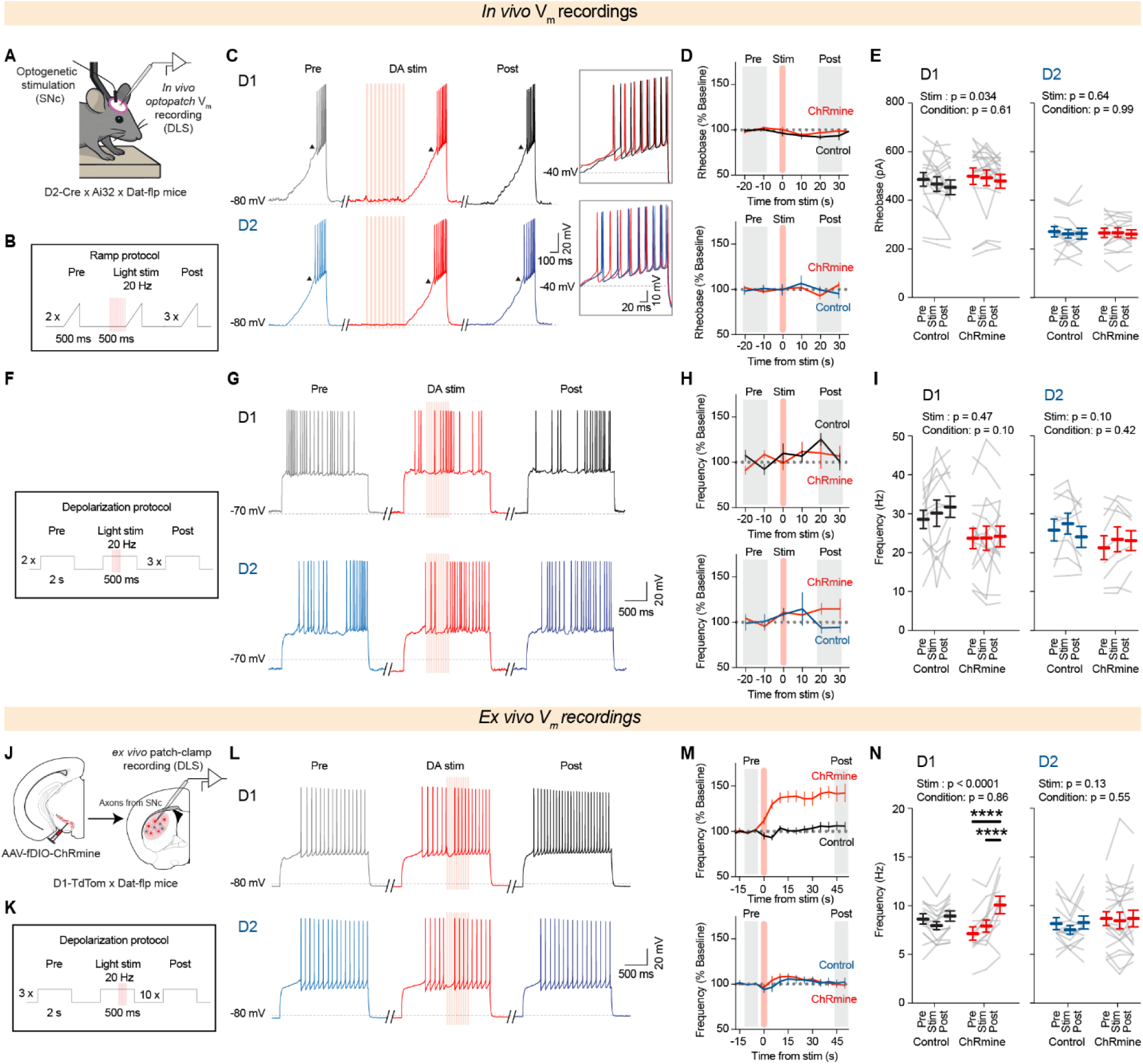
Acute dopamine modulation of D1-SPN excitability is absent *in vivo*. **(A)** Schematic of the *in vivo* recording configuration. AAV-fDIO-ChRmine was expressed in SNc dopamine neurons of D2-Cre x Ai32 x DAT-Flp mice and V_m_ recordings were performed in the DLS of head-fixed awake mice. **(B)** Schematic of the ramp protocol. Two baseline ramps were delivered, followed by a ramp paired with a 500 ms light pulse at 20 Hz, and three post-stimulation ramps with a 10-second inter-ramp interval. **(C)** Representative *in vivo* current ramp traces in D1-SPNs (top) and D2-SPNs (bottom) before, during, and after dopamine release. Insets show first action potentials. **(D)** Rheobase expressed as percentage of baseline over time for D1-SPNs (top) and D2-SPNs (bottom) in ChRmine and control mice. The stimulation ramp is indicated by the pink shaded area. **(E)** Rheobase before, during, and after dopamine release in D1- and D2-SPNs for control and ChRmine mice (repeated measures two-way ANOVA, D1: stim factor: *P* = 0.037, condition: *P* =0.61; Control n = 17 cells from 10 mice, ChRmine n = 19 cells from 14 mice; D2: stim factor: *P* = 0.64, condition: *P* = 0.99, Control n = 15 cells from 8 mice, ChRmine n = 13 cells from 11 mice). **(F)** Schematic of the depolarization protocol. Two baseline 2-second current steps were delivered, followed by a step paired with a 500 ms light pulse at 20 Hz, and three post-stimulation steps with a 10-second inter-step interval. **(G)** Representative *in vivo* current step traces in D1-SPNs (top) and D2-SPNs (bottom) before, during, and after dopamine release. **(H)** Action potential frequency expressed as percentage of baseline over time for D1-SPNs (top) and D2-SPNs (bottom) in ChRmine and control mice. The stimulation step is indicated by the pink shaded area. **(I)** Action potential frequency before, during, and after dopamine release in D1- and D2-SPNs for control and ChRmine mice (repeated measures two-way ANOVA, D1: stim factor: *P* = 0.47, condition: *P* = 0.10; Control n = 12 cells from 7 mice, ChRmine n = 15 cells from 12 mice; D2: stim factor: *P* = 0.42, condition: *P* = 0.10, Control n = 9 cells from 7 mice, ChRmine n = 9 cells from 7 mice). **(J)** Schematic of the ex vivo recording configuration. AAV-fDIO-ChRmine was expressed in SNc dopamine neurons of D1-TdTom x DAT-Flp mice and patch-clamp recordings were performed in DLS slices. **(K)** Schematic of the *ex vivo* depolarization protocol. Three baseline 2-second current steps were delivered, followed by a step paired with a 500 ms light pulse at 20 Hz, and ten post-stimulation steps. **(L)** Representative *ex vivo* current step traces in D1-SPNs (top) and D2-SPNs (bottom) before, during, and after dopamine release. **(M)** Action potential frequency expressed as percentage of baseline over time for D1-SPNs (top) and D2-SPNs (bottom) in ChRmine and control mice in slice. **(N)** Action potential frequency before, during, and after dopamine release in D1- and D2-SPNs for control and ChRmine mice in slice (repeated measures two-way ANOVA, D1: stim factor: *P* < 0.0001, condition: *P* = 0.78; Control n = 14 cells from 4 mice, ChRmine n = 11 cells from 6 mice; D2: stim factor: *P* = 0.13, condition: *P* = 0.55, Control n = 14 cells from 4 mice, ChRmine n = 19 cells from 5 mice). Data are shown as mean ± SEM. Each line represents an individual neuron. \*\*\*\**P* < 0.0001.

Because dopamine has also been reported to enhance excitability during sustained depolarization (*15*), we next delivered dopamine stimulation during a two-second depolarizing current step while monitoring action potential firing (Fig. 2, F and G). Dopamine release had no detectable effect on firing frequency either during stimulation or over the subsequent tens of seconds in either D1- or D2-SPNs (Fig. 2, H and I).

To verify that our experimental approach could detect dopamine-dependent changes in excitability when present, we performed complementary V_m_ recordings in acute brain slices from D1-tdTomato × DAT-Flp mice expressing ChRmine in SNc dopamine neurons (Fig. 2J; fig. S4, A to F). Using the same whole-cell recording configuration, including pipettes of comparable resistance, together with the same depolarizing step protocol paired with dopamine neuron stimulation (Fig. 2, K and L), dopamine release produced a robust and prolonged increase in action potential firing selectively in D1-SPNs (Fig. 2, M and N; Fig. S4, G to K; Table S6), consistent with previous reports (*15*). No effect was observed in D2-SPNs. Thus, the molecular machinery through which dopamine enhances D1-SPN excitability is readily detected *ex vivo* but is not engaged by physiological dopamine transients in the intact brain.

### An S1-guided learning task aligns dopamine release with S1 corticostriatal inputs

Having established an *in vivo* approach for studying identified corticostriatal synapses and finding little evidence that acute dopamine alters the intrinsic physiology of SPNs, we next asked whether dopamine instead shapes corticostriatal circuits during learning. To do so, we developed an optogenetically guided reward task that engaged the same S1→ DLS pathway used to isolate corticostriatal synapses in our electrophysiological experiments. Head-fixed mice learned to lick in response to optogenetic activation of S1 inputs to the dorsolateral striatum to receive a water reward (Fig. 3, A to C). Across days, mice improved, reaching at least 80% correct responses (hit) with decreased false alarm rates below 40% on catch trials (Fig. 3D). Performance was driven by cortical stimulation and not light itself, as shown by a drop in performance when the light source was shifted away from S1 (Fig. 3E; Table S7). Expert mice showed an increase in d’ alongside faster and more stereotyped licking responses (Fig. 3, F to H).

**Fig. 3.**
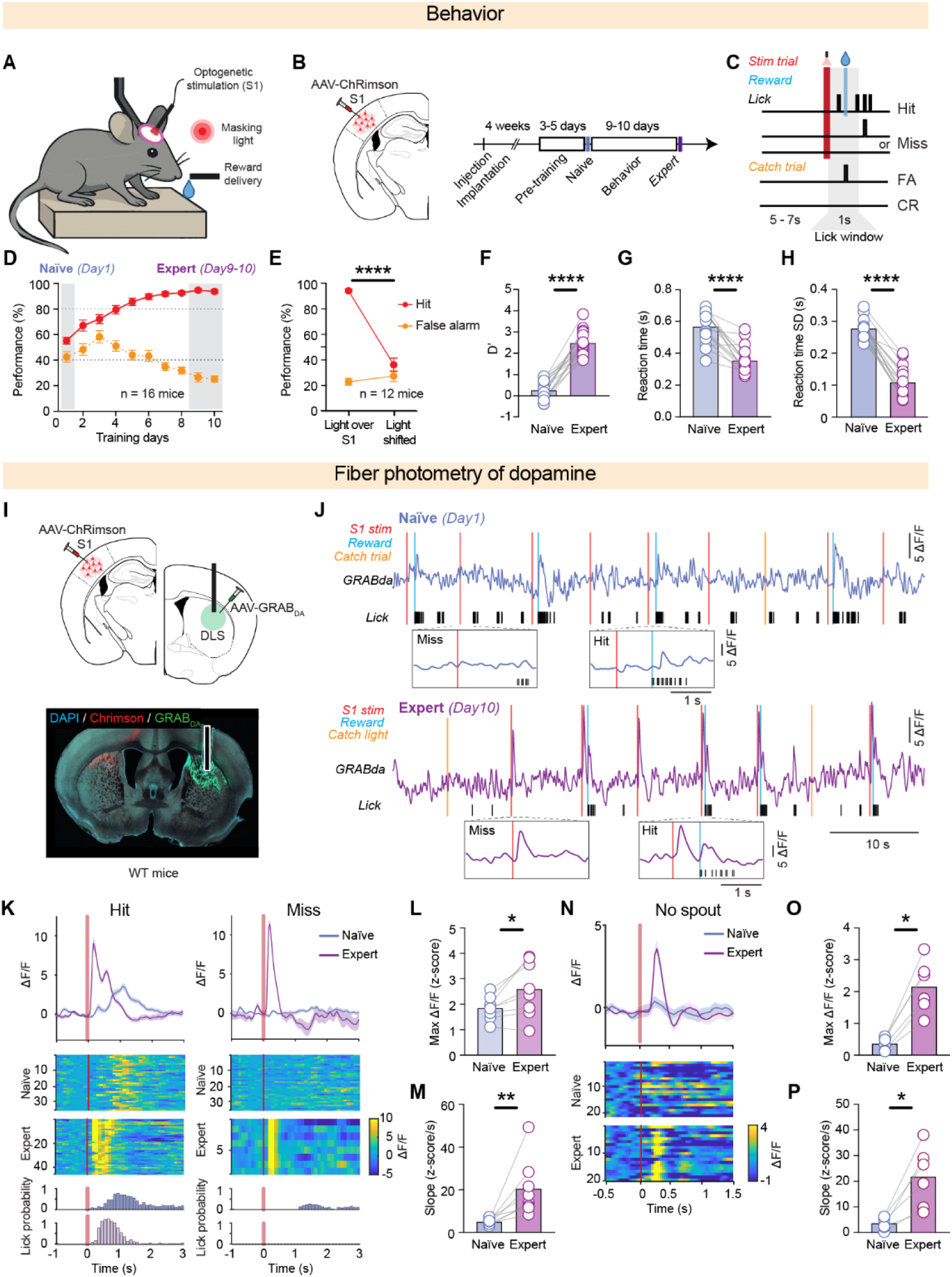
S1-guided learning aligns dopamine release with corticostriatal inputs. **(A)** Schematic of the behavioral task. Head-fixed mice were trained to lick in response to S1 optogenetic stimulation to receive a water reward. A masking light was present in the box to prevent visual detection of the stimulation. **(B)** Experimental timeline. AAV-ChRimson was injected in S1 followed by implantation. 4 weeks later mice underwent pre-training for 3-5 days, and behavioral training for 9-10 days. **(C)** Trial structure. Stimulation trials consisted of an S1 5ms-light pulse paired with a 1-second lick window to get a reward. Catch trials contained no S1 stimulation. Hit: lick on stimulation trial. Miss: no lick on stimulation trial. False alarm (FA): lick on catch trial. Correct rejection (CR): no lick on catch trial. **(D)** Performance across training days showing hit rate and false alarm rate for naïve and expert sessions (n = 16 mice). **(E)** Performance comparison between light delivered over S1 and light shifted away from S1 in expert mice, confirming that performance depends on cortical stimulation and not the light itself (RM Two-way ANOVA, location factor: *P* < 0.0001; n = 12 mice). **(F)** Discriminability index, d’, was significantly higher in expert vs. naïve mice (paired T-test; *P* < 0.0001; n = 16 mice). **(G)** Reaction time was significantly lower in expert vs. naïve mice (paired T-test; *P* < 0.0001; n = 16 mice). **(H)** Reaction time standard deviation (SD) was significantly lower in expert vs. naïve mice (paired T-test; *P* < 0.0001; n = 16 mice). **(I)** Schematic of the fiber photometry configuration. AAV-ChRimson was expressed in S1 and GRAB-DA was expressed in the contralateral DLS of WT mice to monitor dopamine release during behavior. **(J)** Representative fiber photometry traces of GRAB-DA dopamine signals in the DLS during the behavioral task on day 1 (naïve, top) and day 10 (expert, bottom). S1 stimulation times are indicated by red ticks, reward delivery by blue ticks, catch light by yellow ticks, and lick times by black ticks. Representative miss and hit trials are shown at expanded timescale. **(K)** Representative fiber photometry traces of dopamine release aligned to S1 stimulation, hit (left) and miss (right) trials on day 1 and day 10 from the same mouse. Heatmaps show individual trial dopamine signals. Lick activity is shown below. **(L)** Peak dopamine release amplitude during hit trials was significantly higher in expert vs. naïve mice (paired T-test, *P* = 0.002; n = 8 mice). **(M)** Slope of dopamine signal was significantly higher in expert vs. naïve mice (Wilcoxon matched-pairs, *P* = 0.008; n = 8 mice). **(N)** Representative fiber photometry traces of dopamine release in the DLS aligned to S1 stimulation in the absence of the licking spout in naïve and expert mice, showing the emergence of a dopamine response to the predictive cue in expert animals. **(O)** Peak dopamine release amplitude in response to S1 stimulation in absence of spout was significantly higher in expert vs. naïve mice (Wilcoxon matched-pairs, *P* = 0.031; n = 6 mice). **(P)** Slope of dopamine releases amplitude in response to S1 stimulation in absence of spout was significantly higher in expert vs. naïve mice (Wilcoxon matched-pairs, *P* = 0.031; n = 6 mice). Data are shown as mean ± SEM. Each open circle represents an individual mouse fiber \**P* < 0.05, \*\*\**P* < 0.001, \*\*\*\**P* < 0.0001.

We next asked how dopamine signaling evolves as animals learn this S1-guided task. Using fiber photometry of GRABDA2m in the DLS (Fig. 3I), we monitored dopamine dynamics throughout learning. In naïve mice, dopamine transients occurred primarily following reward delivery (Fig. 3, J and K). As learning progressed, dopamine release became biphasic, with a prominent early component emerging immediately following S1 stimulation (Fig. 3, J to M). To determine whether this early signal reflected reward prediction rather than reward consumption, we omitted access to the reward spout on a subset of trials after learning. Under these conditions, robust cue-evoked dopamine release persisted despite the absence of reward consumption (Fig. 3, N to P), demonstrating that dopamine release had become aligned with the reward-predictive S1 input. Consistent with this interpretation, cue-evoked dopamine responses were selectively enhanced on successful (Hit) trials and persisted in the absence of reward, whereas responses on Miss and False Alarm trials remained small throughout learning (Fig. S5). Together, these experiments establish a behavioral paradigm in which dopamine signaling becomes temporally aligned with S1-evoked corticostriatal activity, providing a framework for examining dopamine-dependent plasticity at those synapses.

### Learning strengthens corticostriatal synapses *in vivo*

Previous work has demonstrated that learning can modify corticostriatal synaptic transmission *in vivo* (*29*). If dopamine primarily shapes striatal circuits through learning-dependent synaptic plasticity, such changes should be evident at corticostriatal synapses following learning. We therefore trained naïve and expert mice on the S1-guided licking task (Fig. 4A). To first determine whether learning in this task requires corticostriatal plasticity, we selectively removed NMDA receptors from DLS neurons by injecting AAV-Cre into the DLS of NR1-loxP mice, thereby deleting the obligatory NMDA receptor subunit NR1 locally within the striatum (Fig. 4B, Fig. S6). NR1 knockout mice exhibited normal locomotor activity in an open-field assay and normal baseline licking during pretraining, arguing against a gross motor deficit (Fig. S6). Nevertheless, they showed significantly impaired learning, with reduced *d*′ and increased SD reaction times (Fig. 4, C to E), indicating that successful learning in this task requires intact plasticity within the engaged corticostriatal circuit.

**Fig. 4.**
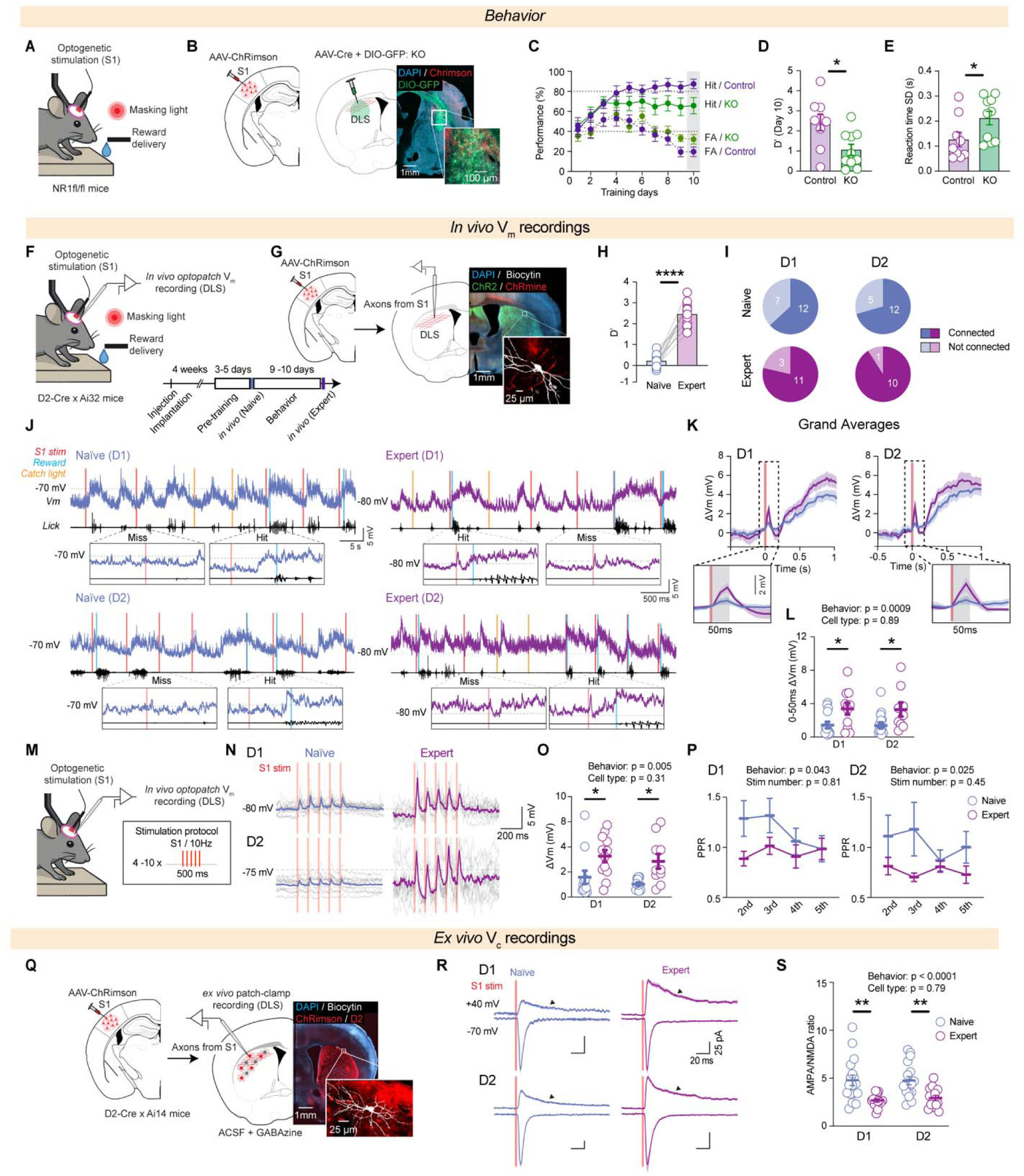
Learning strengthens corticostriatal synapses through NMDAR-dependent plasticity. **(A)** Schematic of the NR1 knockout (NR1KO) behavior configuration. AAV-Cre and AAV-DIO-GFP were injected in the DLS of NR1KO-Cre mice to drive NMDA receptor deletion. **(B)** Left: schematic showing injection of AAV-ChRimson in S1. Right Schematic and representative histology showing GFP expression in DLS and in the inset AAV-ChRimson expressing S1 axons and GFP positive neurons in the DLS. **(C)** Performance across training days in NR1KO (green) and control mice (purple) showing hit rate and false alarm rate. **(D)** Discriminability index d’ in expert mice at day 10 of training was significantly lower in NR1KO vs. control mice (T-test, *P* = 0.012 ; control n = 9 mice, KO n = 10 mice). **(E)** Reaction time standard deviation at day 10 of training was significantly higher in NR1KO vs. control mice (T-test, *P* = 0.045 ; control n = 9 mice, KO n = 10 mice). **(F)** Schematic of the recording configuration. AAV-ChRimson was expressed in S1 of D2-Cre x Ai32 mice and V_m_ recordings were performed in the DLS of head-fixed awake mice during the behavioral task. Experimental timeline showing injection, implantation, pre-training and behavioral training phases. **(G)** Left: schematic showing injection of AAV-ChRimson in S1. Right Schematic and representative histology showing ChR2-YFP expression in DLS and in the inset AAV-ChRimson expressing S1 axons in the DLS and a biocytin-filled neuron recorded *in vivo*. **(H)** Performance across training days in this cohort of mice showing significantly higher D’ in expert vs. naïve mice (paired T-test, *P* < 0.0001; n = 12 mice). **(I)** Proportion of D1-SPNs and D2-SPNs receiving monosynaptic S1 input in naïve and expert mice (D1: naïve n = 12 connected, n = 7 not connected, expert n = 11 connected, n = 3 not connected; D2: naïve n = 12 connected, n = 5 not connected, expert n = 10 connected. n = 1 not connected). **(J)** Representative *in vivo* V_m_ recordings from D1-SPNs (top) and D2-SPNs (bottom) in naïve (blue) and expert (purple) mice during hit and miss trials. S1 stimulation times are indicated by red ticks, reward by light blue ticks and licks (as measured by a piezo film) in black trace. **(K)** Grand average EPSP responses to S1 stimulation in D1-SPNs (left) and D2-SPNs (right) in naïve and expert mice. Shaded area represents SEM. Insets show the onset of the response (first 150 ms), with the dark shaded region indicating the 50 ms window used for the quantification in (L) (D1: naïve n = 14 cells from 10 mice; expert n = 11 cells from 9 mice; D2: naïve n = 16 cells from 11 mice; expert n = 9 cells from 6 mice). **(L)** Maximum depolarization amplitude in the first 50ms after stimulation in D1- and D2-SPNs in naive and expert mice (repeated measures two-way ANOVA, behavior: *P* = 0.0009, cell type: *P* = 0.89; D1: naive n = 14 cells from 10 mice; expert n = 11 cells from 9 mice; D2: naive n = 16 cells from 11 mice; expert n = 9 cells from 6 mice). **(M)** Schematic of the stimulation configuration. In naïve and expert mice, S1 was stimulated at 10 Hz in the absence of the licking spout to isolate synaptic changes from movement-related artifacts. Stimulation protocol: 500ms stimulation at 10 Hz repeated 4-10 times. **(N)** Representative *in vivo* traces from D1-SPNs (top) and D2-SPNs (bottom) in naïve (blue) and expert (purple) mice during 10 Hz S1 stimulation. Scale bars, 200 ms / 2 mV. **(O)** EPSP amplitude of the first peak during 10Hz stimulation in D1- and D2-SPNs from naïve (blue circles) and expert (purple circles) mice in connected cells (two-way ANOVA, behavior: *P* = 0.0005, cell type: *P* = 0.31; naïve: ; D1: naïve n = 12 cells from 9 mice; expert n = 12 cells from 10 mice; D2: naïve n = 11 cells from 7 mice; expert n = 10 cells from 8 mice). **(P)** Paired-pulse ratio during 10Hz stimulation in D1- and D2-SPNs from naïve and expert mice (repeated measures two-way ANOVA; D1: behavior: *P* = 0.043, stimulation: *P* = 0.81; D2: behavior: *P* = 0.025, stimulation: *P* = 0.45). **(Q)** Schematic of the ex vivo voltage clamp (V_c_) recording configuration. AAV-ChRimson was expressed in S1 of D2-Cre x Ai14 mice and patch-clamp recordings were performed in DLS slices in the presence of GABAzine. Representative histology showing ChRimson-expressing S1 axons in the DLS and a biocytin-filled recorded neuron. **(R)** Representative voltage-clamp traces showing AMPA responses at -70 mV and NMDA responses at +40 mV in D1-SPNs (top) and D2-SPNs (bottom) from naïve and expert mice. Asterisks indicate NMDA measurement window at 50 ms after stimulus onset. **(S)** AMPA/NMDA ratio in D1- and D2-SPNs from naïve and expert mice (two-way ANOVA, behavior: *P* < 0.0001, cell type: *P* = 0.79; D1: naïve n = 18 cells from 8 mice; expert n = 13 cells from 7 mice; D2: naïve n = 17 cells from 7 mice; expert n = 14 cells from 6 mice). Data are shown as mean ± SEM. Each open circle represents an individual mouse for D,E and H. Open circle represents an individual neuron for the rest of the figure. * *P* < 0.05, ** *P* < 0.01, **** *P* < 0.0001.

We next examined the synaptic consequences of learning *in vivo*. V_m_ recordings were obtained from D1- and D2-SPNs in naïve and expert mice (Fig. 4 and G), with expert mice exhibiting significantly higher d′ values than naïve mice (Fig. 4H; Table S8). In addition, both cell types received monosynaptic S1 input under both conditions (Fig. 4I), with EPSPs reaching peak amplitude after 21.5 ± 1.4 ms in D1-SPNs and 24.2 ± 1.3 ms in D2-SPNs. Intrinsic membrane properties were unchanged following learning (Fig. S6; Table S9). Early EPSP responses, measured within the first 50 ms of cortical stimulation, were significantly larger in expert than naïve mice during hit trials in both D1- and D2-SPNs (Fig. 4, K to L). To further assess synaptic strength independently of ongoing behavior, we delivered trains of S1 stimulation (10 Hz) in the absence of the licking spout (Fig. 4M). Expert mice again exhibited larger EPSP amplitudes (Fig. 4, N and O), accompanied by a reduction in paired-pulse ratio (Fig. 4P) in both cell types.

Finally, to determine whether learning was associated with additional synaptic signatures consistent with corticostriatal plasticity, we measured AMPA/NMDA ratios in acute slices prepared from naïve and expert mice (Fig. 4Q). Learning was associated with a significant decrease in the AMPA/NMDA ratio (Fig. 4R and S), consistent with previous observations following corticostriatal learning (*30*).

### Dopamine selectively gates corticostriatal plasticity in D1-SPNs

We next asked whether dopamine signaling is required for the induction of this plasticity. Although behavioral photometry recordings were performed in the contralateral dorsolateral striatum, direct optogenetic activation of S1 reliably evoked dopamine release in the ipsilateral dorsolateral striatum (Fig. 5, A to C), consistent with previous reports (*31*, *32*). The signal was abolished by the D2 receptor antagonist raclopride, as expected for the D2 receptor-based GRABDA2m sensor (Fig. 5C). We therefore used repeated S1 stimulation over eight days to engage endogenous dopamine signaling while driving activity in the same corticostriatal pathway.

**Fig. 5.**
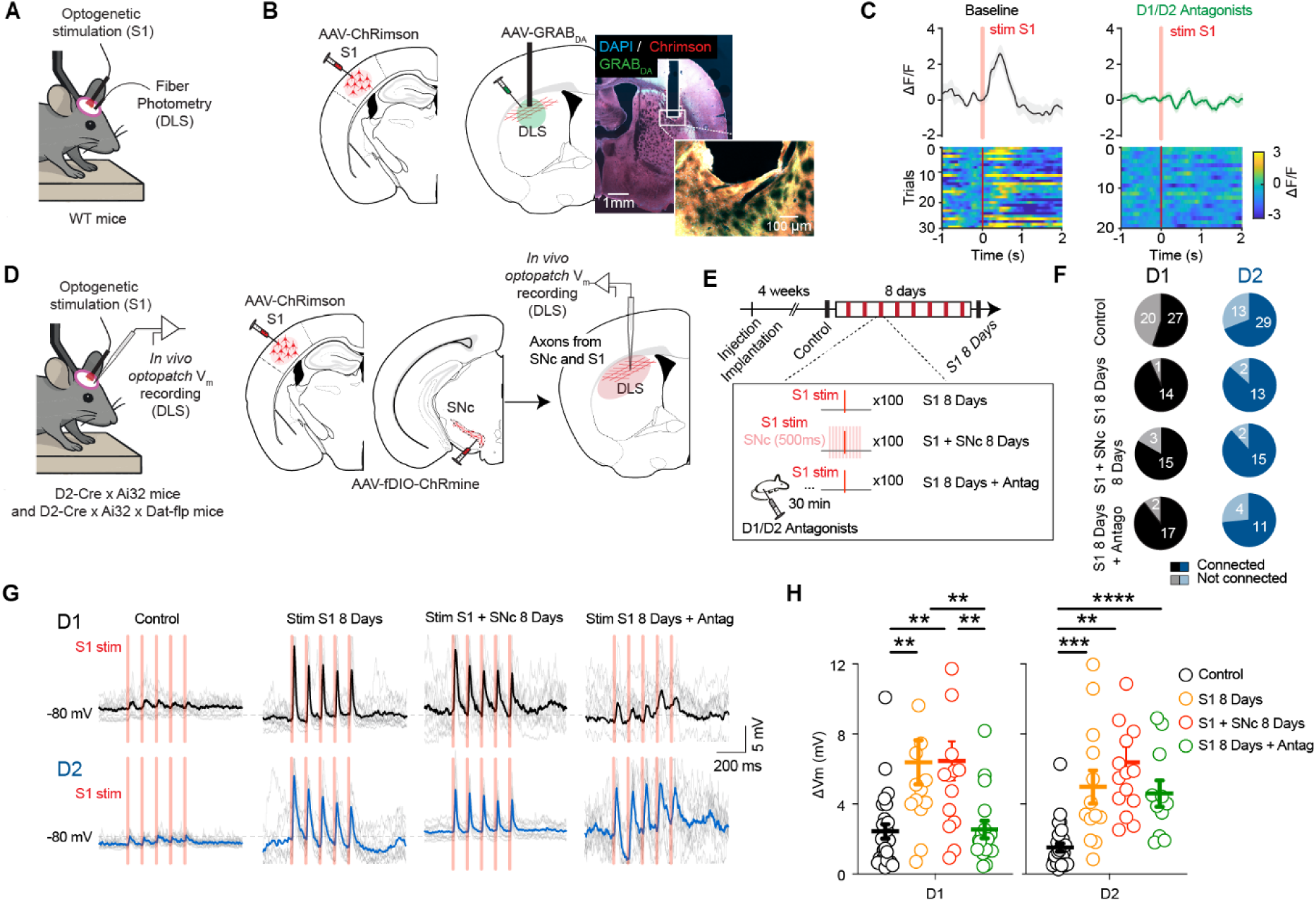
Dopamine selectively gates corticostriatal plasticity in D1-SPNs. **(A)** Schematic of the fiber photometry configuration. AAV-ChRimson was expressed in S1 and GRAB-DA was expressed ipsilaterally in the DLS of WT mice to monitor dopamine release during S1 stimulation. **(B)** Representative histology showing AAV-ChRimson expression in S1 and GRAB-DA expression in the DLS with fiber optic placement. **(C)** Representative fiber photometry traces (top) and trial heatmaps (bottom) of dopamine release in the DLS aligned to S1 stimulation. Left: at baseline Right: after D1/D2 receptor antagonist injection. **(D)** Schematic of the *in vivo* recording configuration. AAV-ChRimson was expressed in S1 and AAV-fDIO-ChRmine was expressed in SNc dopamine neurons of D2-Cre x Ai32 x DAT-Flp mice, allowing independent co-stimulation of cortical inputs and dopamine neurons in the S1 + SNc 8 days conditions. V_m_ recordings were performed in the DLS. **(E)** Experimental timeline showing injection in S1 and implantation followed by four conditions: “control”: mice received no S1 stimulation; “S1 8 days”: mice received 100 5ms S1 stimulations per day for 8 days; “S1 + SNc 8 days”: mice received 100 paired 5ms S1 and 500ms 20Hz SNc stimulations per day for 8 and “S1 8 days + antagonist”: mice received the D2 antagonist raclopride and D1 antagonist SCH 23390 30 minutes before each daily stimulation session of 100 5 ms S1 stimulations for 8 days. In this last condition, *in vivo* V_m_ recordings commenced the following day in the absence of any drug injection. **(F)** Proportion of D1-SPNs (left) and D2-SPNs (right) receiving monosynaptic S1 input across the four conditions: control, S1 8 days, and S1 8 days with antagonists (D1: control: n = 27 connected, n = 22 not connected; S1 8 days: n = 14 connected, n = 1 not connected; S1 + SNc 8 days: n = 15 connected, n = 3 not connected; S1 8 days + antagonist : n = 17 connected, n = 2 not connected; D2 control: n = 29 connected, n = 13 not connected; S1 8 days D2: n = 13 connected, n = 2 not connected; S1 + SNc 8 days : n = 15 connected, n = 2 not connected; S1 8 days + antagonist D2: n = 11 connected, n = 4 not connected). **(G)** Representative *in vivo* traces from D1-SPNs (top) and D2-SPNs (bottom) in control, S1 stim 8 days, S1 stim + SNc stim 8 days and S1 stim 8 days with antagonist conditions during S1 stimulation. Individual trials shown in gray, average in color. **(H)** Peak EPSP amplitude in D1- and D2-SPNs across control and the four conditions (One-way ANOVA, D1: *P* < 0.0001, control: n = 27 cells from 17 mice; S1 8 days n = 14 cells from 5 mice; S1 + SNc 8 days n = 15 cells from 7 mice; S1 8 days + antagonist n = 17 cells from 8 mice; D2: *P* < 0.0001; control: n = 29 cells from 17 mice; S1 8 days n = 13 cells from 5 mice; S1 + SNc 8 days n = 15 cells from 10 mice; S1 8 days + antagonist n = 11 cells from 6 mice). Data are shown as mean ± SEM. Each open circle represents an individual neuron. (\*\**P* < 0.01, \*\*\**P* < 0.001, \*\*\*\**P* < 0.0001.

To determine whether dopamine signaling is required for the induction of corticostriatal plasticity, we compared four experimental groups: unstimulated controls, mice receiving repeated S1 stimulation over eight days, mice receiving S1 stimulation paired with optogenetic activation of SNc dopamine neurons, and mice receiving S1 stimulation together with systemic D1 and D2 receptor antagonists (Fig. 5D and E). Dopamine receptor blockade reduced spontaneous locomotor activity in the open field (Fig. S7, A to C), consistent with the established role of dopamine signaling in movement, and abolished spontaneous GRABDA2m signals (Fig. S7, D to F; Table S11). *In vivo* V_m_ recordings recordings confirmed monosynaptic S1 input onto both D1- and D2-SPNs in all four experimental groups (Fig. 5, E to G).

Repeated S1 stimulation alone was sufficient to produce robust strengthening of identified corticostriatal synapses in both D1- and D2-SPNs relative to unstimulated controls (Fig. 5, G and H; Table S10). Pairing S1 stimulation with simultaneous activation of SNc dopamine neurons throughout the stimulation protocol did not further enhance synaptic strengthening (Fig. 5, G and H), indicating that endogenous dopamine release elicited by cortical activity is sufficient to fully engage the dopamine-dependent component of this plasticity.

Remarkably, however, blocking dopamine receptors throughout the stimulation protocol completely abolished synaptic strengthening in D1-SPNs while leaving plasticity in D2-SPNs intact (Fig. 5, G and H). Thus, although repeated activation of the same corticostriatal pathway strengthens synapses onto both direct- and indirect-pathway neurons, dopamine signaling is selectively required for the induction of D1-SPN plasticity.

## Discussion

Our results identify learning-related corticostriatal plasticity as the principal physiological consequence of dopamine signaling during behavior. Using an *in vivo* preparation that enabled recordings from identified corticostriatal synapses, we found that associative learning strengthens synapses onto both D1- and D2-SPNs, yet dopamine signaling is selectively required for the induction of D1-SPN plasticity. Remarkably, despite this essential role during learning, neither endogenous nor experimentally manipulated phasic dopamine signaling produced detectable changes in membrane potential dynamics or intrinsic excitability. Together, these findings suggest that dopamine primarily shapes striatal circuits by selectively gating learning-related synaptic plasticity rather than by acutely modulating the physiological state of striatal neurons.

We replicated the canonical finding that dopamine increases the excitability of D1-SPNs without affecting D2-SPNs *ex vivo* (*15*), confirming that the cellular machinery through which dopamine can regulate intrinsic membrane properties is present in our hands. Rather than contradicting this literature, our findings suggest that dopamine-dependent changes in excitability are not elicited by physiological phasic dopamine release in the intact brain. In the intact brain, ongoing midbrain activity continuously engages dopamine receptors and downstream signaling pathways, potentially establishing a baseline state of enhanced excitability that occludes the impact of additional dopamine fluctuations (*33*, *34*). Because acute slice recordings are performed after prolonged interruption of endogenous dopamine signaling, dopamine application may partially restore a dopamine-dependent excitability state that is already established in the intact brain. In this framework, D1-SPNs *in vivo* may operate near a dopamine-dependent excitability set point maintained by ongoing tonic dopamine signaling, such that physiological phasic increases in dopamine produce little additional effect on membrane potential dynamics or firing threshold. Thus, the excitability changes observed in slice preparations and the lack of acute modulation *in vivo* may reflect complementary manifestations of dopamine action under distinct physiological conditions.

Repeated activation of a corticostriatal pathway drove robust synaptic strengthening in both D1- and D2-SPNs. Although previous work has established distinct molecular requirements for corticostriatal plasticity in these two cell types (*12*, *35*), whether associative learning strengthens corticostriatal synapses onto both pathways during behavior has remained unknown. Despite exhibiting comparable synaptic strengthening, the underlying mechanisms proved fundamentally different. Blocking D1 and D2 receptors throughout the stimulation protocol selectively abolished D1-SPN strengthening while leaving D2-SPN plasticity intact, demonstrating that learning-related corticostriatal plasticity engages distinct molecular mechanisms in the two striatal output pathways. The dopamine dependence of D1-SPN plasticity is consistent with the established role of D1-cAMP-PKA signaling in corticostriatal potentiation (*9*, *13*). By contrast, the persistence of D2-SPN strengthening in the absence of dopamine receptor signaling suggests the involvement of adenosine A2A receptor signaling, which has been shown to support LTP in indirect-pathway SPNs (*12*, *36*, *37*). Cholinergic interneurons may also contribute, as coordinated cholinergic pauses and dopamine transients have been implicated in corticostriatal plasticity and reinforcement learning *in vivo* (*21*, *38*, *39*). Finally, pairing S1 stimulation with optogenetic activation of SNc dopamine neurons failed to further enhance synaptic strengthening, suggesting that repeated S1 activation recruits sufficient endogenous dopamine to fully engage D1-dependent plasticity under these learning conditions.

Finally, our findings raise an interesting question about hyperdopaminergic states. If physiological phasic dopamine is insufficient to drive acute changes in striatal physiology, what happens when dopamine is massively and persistently elevated, for example during cocaine exposure? Our observation that co-stimulating SNc dopamine neurons alongside cortical inputs does not enhance plasticity beyond cortical stimulation alone suggests that the physiological plasticity mechanism described here saturates at relatively low dopamine levels. This interpretation is consistent with recent behavioral studies showing that physiological dopamine transients exert surprisingly limited effects on ongoing movement and vigor, whereas supraphysiological dopamine stimulation can produce robust behavioral changes. Together, these findings raise the possibility that once dopamine signaling exceeds its normal physiological range, qualitatively different circuit mechanisms may become engaged. Such a framework may help explain why chronic cocaine exposure produces profound structural and synaptic remodeling (*40–42*) that appears disproportionate to the role of dopamine during normal learning. Cocaine, by blocking the dopamine transporter (*43*), sustains supraphysiological dopamine levels far beyond those generated by natural rewards (*41*, *44*). Whether addiction reflects an extension of the dopamine-dependent plasticity described here beyond its normal physiological operating range, or instead recruits fundamentally distinct plasticity mechanisms, remains an important question for future work.

## Acknowledgments

We thank all Sippy lab members for their time and assistance, especially Corryn Chaimowitz and Sulekh Fernando-Peiris for mouse colony maintenance and technical assistance and Jan Klee for help with development of the behavior scripts and analysis. We thank Vincent Robert for homemade analysis code for *ex vivo* electrophysiology and behavior, and for feedback on the manuscript. We are grateful to Christine Constantinople and Michael long for helpful comments on the manuscript.

## Author contributions

MD, NXT and TS conceived the project and designed experiments; MD performed all the experiments and analysis; YCY assisted with stereotaxic injections, fiber photometry recording, behavior, data organization and analysis; MD, NXT and TS wrote the paper; NXT and TS acquired funding and supervised personnel; TS provided resources and managed the project.

## Data and materials availability

All data are available in the manuscript, or the supplementary materials will be deposed at Zenodo. All original code will be available at Zenodo. Further information and requests for materials and additional details required to reanalyze the data reported here should be directed to and will be fulfilled by the corresponding author.

## Methods

### Animals

All experiments were conducted with 8–18-week-old male and female mice in accordance with protocols approved by the NYU Langone Health Institutional Animal Care and Use Committee (protocol #PROTO201900059). D2-Cre BAC transgenic mice were obtained from GENSAT and purchased through the Mutant Mouse Regional Resource Centers (MMRRC: 32108). DAT-flp mice (JAX: 035436) were used for optogenetic activation and inhibition of dopamine neurons. For *in vivo* whole-cell recordings with optogenetic cell-type identification, D2-Cre mice were crossed with Ai32 mice (JAX: 024109) to generate D2-Cre × Ai32 offspring. For *ex vivo* slice experiments, both Drd1a-tdTomato mice (JAX: 016204) and D2-Cre mice crossed with Ai14 (JAX: 007914) were used. For conditional deletion of the NR1 subunit of NMDA receptors in the DLS, NR1 homozygous floxed mice (NR1fl/fl; JAX:036352) were used. All mice were bred on a C57Bl/6J background and housed in groups on a reverse light/dark cycle (lights on 11 pm to 11 am) at 22 ± 2°C with food available ad libitum.

### Surgical procedures and viral injections

All surgeries were performed under isoflurane anesthesia (1.5–2%). Body temperature was maintained with a closed-loop heating pad throughout. A lightweight metal head post was implanted on the skull at least 4 weeks before recording. Viral injections were performed during the same surgery as head post implantation. A glass pipette was lowered to the target site, and 300 nl of virus was injected at a flow rate of 5 nl/s using a Drummond Nanoject III. The pipette was left in place for at least 10 minutes after injection before slow retraction to prevent backflow. For fiber photometry experiments, an optical fiber (400 µm diameter, NA = 0.5; RWD Life Science) was implanted above the DLS immediately after virus injection and secured with dental cement.

For optogenetic stimulation, an optical fiber was implanted above the SNc. A small craniotomy for *in vivo* whole-cell recordings was made acutely on the day of recording session.

For optogenetic activation of dopamine neurons, DAT-Flp mice received unilateral injections of pAAV-Ef1a-fDIO-ChRmine-mScarlet (ZI Virology; 5.20×10¹² vg/ml, diluted 1:2; 300 nl) into the SNc (3.1 mm posterior, 1.6 mm lateral, 4.2–3.8 mm ventral to bregma). For optogenetic silencing of dopamine neurons, DAT-Flp mice received unilateral injections of AAV-EF1a-FRT-FLEX-GtACR2-EYFP-Kv2.1C-linker-TlcnC (ZI Virology; 8.20×10¹² vg/ml, diluted 1:2; 300 nl) into the SNc at the same coordinates. For S1 optogenetic stimulation, AAV1.hSyn.ChrimsonR.tdTomato (UNC/Boyden; 3.7×10¹² vg/ml, diluted 1:4; 300 nl) was injected into S1 (1.7 mm posterior, 3.0 mm lateral, 0.6–0.4 mm ventral to bregma).

For dopamine fiber photometry dopamine neuron manipulation, mice received injections of either pAAV-hSyn-GRAB_DA2m (Addgene #140553-AAV9; 2.4×10¹³ GC/ml, diluted 1:4; 300 nl) or pAAV-hSyn-GRAB_rDA3m (BrainVTA #PT4746; 5.65×10¹2 GC/ml, diluted 1:3; 300 nl) into the DLS at the coordinates above. For conditional deletion of the NR1 subunit of NMDA, NR1 homozygous floxed mice received co-injections of AAV1.hSyn.Cre (Addgene #105553-AAV1; 2.4×10¹³ GC/ml, diluted 1:3; 500 nl) and a Cre-dependent GFP reporter (AAV1.CAG.FLEX.eGFP; UNC/Boyden; 4.4×10¹² vg/ml, diluted 1:3; 500 nl) into the DLS. Littermate controls from the same NR1fl/fl background received injections of the non-Cre-dependent GFP virus alone at the same coordinates and concentration (AAV1.CAG.eGFP; UNC/Boyden; 4.4×10¹² vg/ml, diluted 1:5; 500 nl).

### *In vivo* membrane potential recordings

Mice were habituated to head fixation over 3 days in increasing time increments prior to recording. On the day of recording, a small craniotomy was made under isoflurane anesthesia over the DLS at the coordinates described above. Mice were allowed to recover from anesthesia for 1 hour before being transferred to the recording setup.

Whole-cell patch-clamp recordings were obtained in current-clamp mode using 6–8 MΩ borosilicate glass pipettes filled with an internal solution containing (in mM): 135 K-methylsulfonate, 5 KCl, 0.1 EGTA-KOH, 10 HEPES, 2 NaCl, 5 MgATP, 0.4 Na GTP, 10 Na -phosphocreatine, with 2–4 mg/ml biocytin added for post-hoc morphological identification. Membrane potential was not corrected for liquid junction potential. Signals were amplified using a Multiclamp 700B amplifier (Axon Instruments), digitized at 20 kHz using a National Instruments acquisition board (BNC-2110), and recorded using Wavesurfer software (HHMI Janelia Research Campus).

SPNs were identified as previously described (*45*, *46*), based on their characteristic hyperpolarized resting membrane potential, low spontaneous firing rate, low input resistance, and high rheobase. At the start of each recording, intrinsic properties were assessed by injecting a series of current steps from -200 pA in increments of +25 pA. Recordings were included only if the neuron exhibited a stable resting membrane potential and overshooting action potentials. D1 and D2 SPNs were distinguished using the Optopatcher (A-M Systems), which allows simultaneous optical stimulation and whole-cell recording through the same patch pipette. In D2-Cre × Ai32 mice, 500 ms pulses of 470 nm light (1–2 mW at the fiber tip) were delivered through the pipette to identify ChR2-expressing D2 SPNs by their step-like depolarization and action potential firing (Fig. S2). ChR2-negative neurons showing no response or a small hyperpolarization were classified as putative D1 SPNs.

For optogenetic manipulation of SNc dopamine neurons, light was delivered through the chronically implanted fiber placed above the SNc at the time of viral injection. For dopamine neuron silencing, a continuous light ramp was delivered at 1 mW over 7–10 seconds to activate GtACR2-expressing neurons. For dopamine neuron activation, 500 ms pulse trains at 20 Hz and 2 mW were delivered to activate ChRmine-expressing neurons, except where noted in the supplementary figures, where different stimulation parameters were used.

### *In vivo* data analysis

All *in vivo* data analysis was performed in MATLAB using custom-written algorithms.

A neuron was considered monosynaptically connected to S1 if the trial-averaged Vm response to optogenetic stimulation was at least 0.5 mV depolarization, and if a detectable depolarization was present in at least 50% of individual trials within the 4 to10 trials. Neurons that did not meet both criteria were considered as not connected. Subthreshold membrane potential dynamics were characterized by time-frequency decomposition using complex Morlet wavelet convolution (5 cycles per wavelet, 1–10 Hz). To compare spectral content before and after SNc manipulation, continuous Vm recordings were segmented into non-overlapping 500 ms windows and power spectral density was estimated using the multitaper method (time-bandwidth product of 3). The first and last 5 seconds of each recording were excluded to minimize edge artifacts. PSDs were averaged across windows within each epoch per cell, then across cells within each group, and are presented as mean ± SEM. Summed spectral power in the 1–4 Hz band was used as a measure of low-frequency oscillatory activity. The mean and standard deviation of Vm were computed per 15 seconds before and 15 seconds after the stimulation.

Intrinsic electrophysiological properties including resting membrane potential, input resistance, and rheobase were extracted as previously described (*46*). Input resistance was measured as the slope of the linear fit to voltage responses evoked by hyperpolarizing current steps from -200 pA to 0 pA in increments of 25 pA. Rheobase was defined as the minimum current required to elicit at least one action potential, determined from a series of depolarizing current steps injected at the start of each recording.

For ramp protocol experiments, current ramps were injected at 50–100 pA above the approximate rheobase to reliably elicit 2–3 action potentials at the first ramp stimulation. Cells that failed to fire during any ramp sweep were excluded from all analyses. For depolarizing step experiments, current steps were delivered at 50–70 pA above rheobase to elicit consistent but submaximal firing, and any cell that failed to fire during at least one step was excluded.

For analysis of S1 stimulation-evoked Vm responses, recordings consisted of 5 baseline trials, 5 paired stimulation trials, and 5 post-stimulation trials. Peri-stimulus traces were baseline-subtracted using the mean Vm in the 500 ms window preceding each stimulation onset. Response amplitude was defined as the maximum depolarization from baseline within the first 50 ms following stimulation onset, averaged across trials per cell.

### Fiber photometry

Fiber photometry signals were acquired using a Neurophotometrics FP3002 system (MBF Bioscience), except for simultaneous Vm and dopamine recordings (Fig. S3), for which a Doric fiber photometry system was used with a fluorescence mini-cube (Doric, FMC5_E1(460–490)_F1(500–540)_E2(555–570)_F2(580–680)_S) and a Newport photoreceiver (2151, DC mode). For all experiments, excitation light was delivered through the chronically implanted fiber via a fiber-optic patch cord at 20–40 µW measured at the tip. For green GRABDA2m, a 470 nm LED was used for excitation with emission collected in the 500–540 nm band. For red GRABDA3m, a 560 nm LED was used with emission collected in the 580–680 nm band.

Photometry signals were either acquired at 20kHz and downsampled to 30 Hz or acquired at 30Hz. ΔF/F was computed as (F − F)/F, where F is baseline fluorescence estimated by interpolating the lower envelope of fluorescence values within 30-second sliding windows across the full recording session. For within-mouse comparisons across conditions, ΔF/F traces are shown. For across-mouse quantification and group comparisons, signals were additionally z-scored across the full session. Peri-event traces were extracted in a window of 2 seconds before to 5 seconds after each event of interest, and baseline-subtracted using the 0.5 seconds immediately preceding the event. Response amplitude was quantified as the peak value within 1.5 seconds of event onset, response slope as the maximum rate of rise between 0.1 and 0.4 seconds post-event, and area under the curve (AUC) over a 1-second window following event onset.

### S1 guided learning task

Mice were food-deprived starting 2 days before training onset and maintained at 85% of their initial body weight throughout the experiment. Prior to task training, mice underwent 3–4 days of pre-training in which they were head-fixed in the behavioral box and trained to lick a spout for 10% sucrose water reward delivered by a solenoid valve, with reward available every 2-4 seconds to familiarize mice with the setup.

For the lick task, mice received approximately 100 trials per day consisting of S1 stimulation trials and catch trials. S1 stimulation was delivered as a single 5 ms pulse of red light (625 nm) at 4 mW through the glue over the bone over S1, activating Chrimson-expressing corticostriatal neurons. Mice were required to lick within 1 second of stimulation onset to receive a reward. Catch trials, in which no S1 stimulation was delivered, were interleaved at a proportion starting at approximately 20% and increasing to approximately 50% over the course of training. The behavioral box was maintained under constant red background illumination throughout all training sessions. To confirm that mice responded to the cortical stimulation rather than any light cue, a separate LED positioned above the head was used to deliver a visual distractor during both S1 and catch trials starting around days 3.

Learning was assessed per session and defined as achieving a hit rate of at least 80% and a false alarm rate below 40%. The first day of training was designated as the naïve timepoint. For fiber photometry and *ex vivo* slice experiments, the expert timepoint was defined as day 9 or 10 of training. For NR1KO experiments, mice underwent the same training protocol of 100 stimulation trials per day for 10 days regardless of performance. The experimenter was blind to injection condition throughout training and data collection for all NR1 KO experiments.

Behavioral performance was quantified per session. Hit rate was defined as the proportion of S1 stimulation trials in which the mouse licked within 1 second of stimulation onset. False alarm rate was defined as the proportion of catch trials in which the mouse licked within the same time window. Discriminability was computed as d-prime = z(hit rate) − z(false alarm rate) per session. Reaction time was measured as the latency from S1 stimulation onset to the first lick on hit trials.

Behavioral data acquired during the S1-lick task and video data was recorded using HARP Behavior board and custom workflows using the BONSAI software package and can be accessed on Github.

### Open field recordings

Mice were placed in the center of a 50 × 50 cm open field arena with white Plexiglas flooring and 35 cm walls, illuminated from above at 250 lux, and allowed to explore freely for 1 hour. Locomotor trajectories were tracked using a camera on top, processed with DeepLabCut and analyzed with custom IgorPro code.

### *Ex vivo* electrophysiology

Acute coronal slices (280 µm) were prepared from mice that had completed the training protocol (naïve or expert, as described above). Mice were deeply anesthetized with isoflurane and perfused transcardially with ice-cold oxygenated cutting solution containing (in mM): 110 choline chloride, 2.5 KCl, 25 glucose, 25 NaHCO3, 1.25 NaH2PO4, 0.5 CaCl2, 7 MgCl2, 11.6 L-ascorbic acid and 3.1 sodium pyruvate, as previously described. The brain was rapidly extracted and slices were cut on a vibratome (Leica VT1200S) in the same solution, then transferred to ACSF containing (in mM): 125 NaCl, 2.5 KCl, 25 glucose, 25 NaHCO3, 1.25 NaH2PO4, 2 CaCl2 and 1 MgCl2, continuously bubbled with 95% O2/5% CO2. Slices were transfered at 32°C then stored at room temperature until recording. All recordings were performed at 30–32°C with continuous ACSF perfusion.

For voltage-clamp recordings of corticostriatal synaptic currents, 4–5 MΩ borosilicate glass pipettes were filled with a cesium-based internal solution containing (in mM): 120 Cs-methanesulfonate, 10 CsCl, 10 HEPES, 10 phosphocreatine, 0.2 EGTA, 8 NaCl and 2 MgATP (pH 7.25, adjusted with CsOH). Experiments were performed in the presence of gabazine (10 µM, Hello Bio) to block GABAergic transmission. Corticostriatal synaptic inputs were activated optogenetically by stimulating Chrimson-expressing S1 axons with 595 nm light delivered through a 40× objective. AMPA currents were measured at a holding potential of -70 mV as the peak amplitude of the light-evoked response. NMDA currents were measured at a holding potential of +40 mV, 50 ms after stimulus onset, when AMPA receptor-mediated currents had decayed. The AMPA/NMDA ratio was calculated as AMPA current / NMDA current. The NMDA current measured at +40 mV was confirmed in a subset of cells by bath application of D-APV (50 µM, Hello Bio), which abolished any current after 50 ms.

For experiments assessing dopamine-dependent plasticity, recordings were obtained in current-clamp using the internal solution, optopatch holder and pipettes from the *in vivo* setup. Dopamine release from SNc terminals was evoked by delivering 500 ms trains of 595 nm light pulses at 20 Hz through the 40x objective during a depolarization step.

*Ex vivo* recording were analyzed with custom IgorPro codes.

### Pharmacology

To confirm that dopamine receptor activation was required for S1 stimulation-evoked plasticity in the DLS, dopamine receptor antagonists were administered intraperitoneal prior to fiber photometry recordings. Mice expressing GRABDA2m in the DLS and ChRmine in S1 received injections of the D1 receptor antagonist SCH-23390 (0.2 mg/kg) and the D2 receptor antagonist raclopride (2 mg/kg). Dopamine fluctuations were monitored in baseline with fiber photometry and during S1 stimulation (single 5 ms pulse, 625 nm, 4 mW) before and after injection. Complete abolishment of dopamine transients was confirmed within 30 minutes of injection, after which no further responses to S1 stimulation were detected.

### Histology

All mice were transcardially perfused with 4% paraformaldehyde in phosphate-buffered saline (PBS) at the end of the experiment. Brains were post-fixed for 24 hours in the same solution, then transferred to PBS. Coronal sections were cut at 100 µm on a vibratome (Leica VT1000S) and mounted with DAPI-containing fluorescence mounting medium (Fluoromount-G, Southern Biotech). Images were acquired on a slide scanner (Olympus VS200) with a 20× objective. All mice were included in the final dataset only after histological verification of viral expression and fiber or electrode placement. Animals with mistargeted injections, absent viral expression, or fiber placement outside the DLS were excluded from analysis.

For *in vivo* whole-cell recordings, 100 µm sections were incubated with Streptavidin conjugated to Alexa Fluor 647 (1:2000, Invitrogen) as previously described (*46*). Patched cells were included in the analysis only if biocytin was found within the DLS, especially in the region of Chrimson-expressing S1 axonal projections.

For NR1 KO experiments, Cre-mediated recombination in injected mice was verified by the presence of Cre-dependent GFP expression. Control mice were verified by the presence of non-Cre-dependent GFP expression at the injection site.

To verify dopamine neuron targeting in DAT-Flp mice, tyrosine hydroxylase (TH) immunostaining was performed on SNc. Sections were permeabilized and blocked for 1 hour prior to overnight incubation at 4°C with a rabbit anti-TH primary antibody (1:1000; AB152, Millipore). Sections were then incubated with a donkey anti-rabbit Alexa Fluor 647 secondary antibody (1:1000; Thermo Fisher) for 2 hours at room temperature before mounting.

### Statistics

Data are presented as mean ± SEM. Statistical analyses were performed using Prism 9 (GraphPad) and custom MATLAB code. Normality was assessed using the D’Agostino–Pearson test. For comparisons between two unpaired groups, two-tailed Student’s t-tests or Mann– Whitney tests were used for normally and non-normally distributed data, respectively. For paired comparisons, two-tailed paired t-tests or Wilcoxon signed-rank tests were used accordingly. For multiple comparisons, one-way or two-way ANOVA was followed by Sidak’s or Tukey’s post-hoc test as appropriate. A significance threshold of *P* < 0.05 was applied throughout. *P* values are indicated as follows: * *P* < 0.05; ** *P* < 0.01; *** *P* < 0.001; **** *P* < 0.0001. Error bars and shaded areas around averages represent ± SEM.

