## Supplemental Figures S1-S7 and Tables S1-S11 for "The Cellular and Synaptic Actions of Dopamine During Behavior"

**Supplementary Figures**

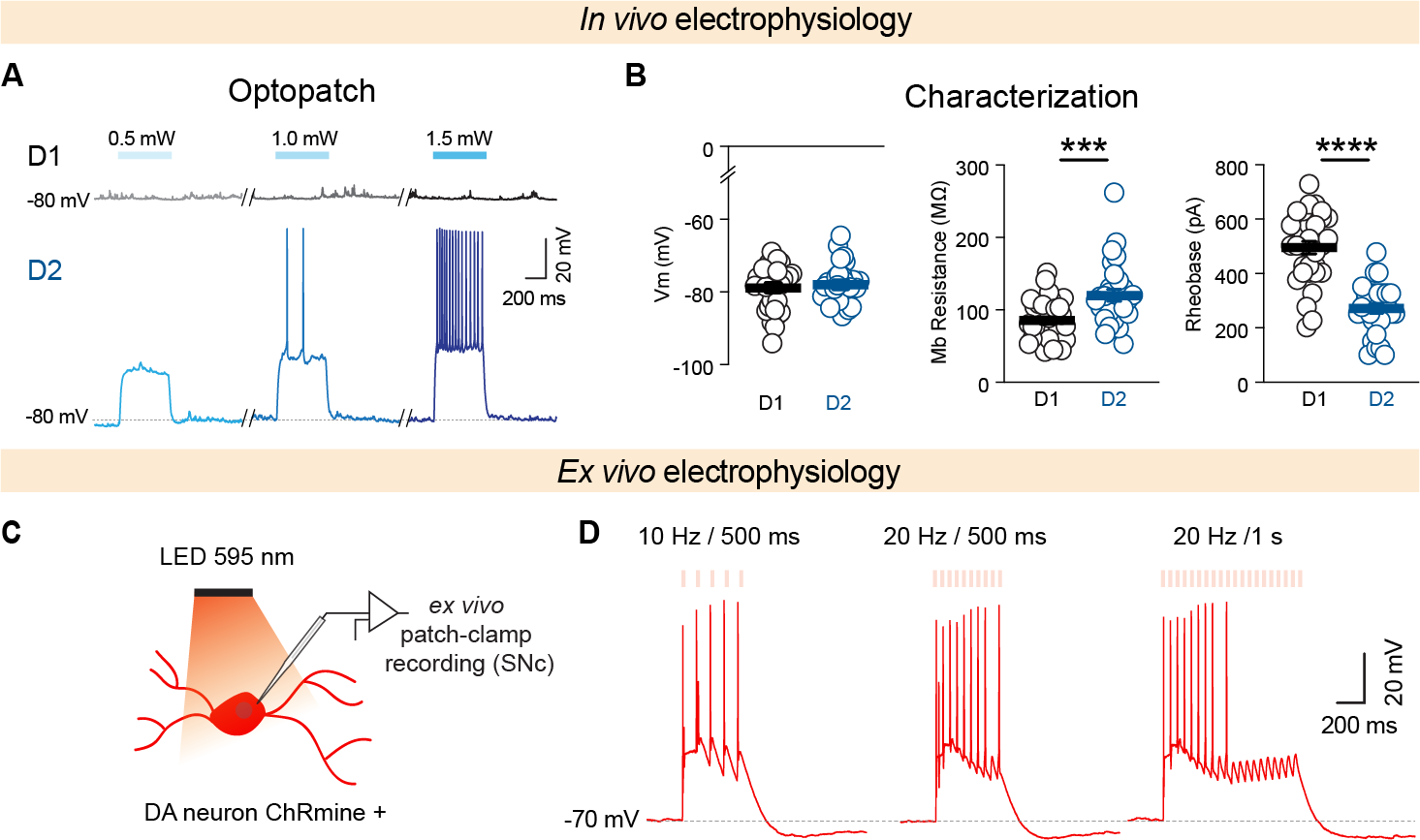

**Fig. S1. Validation of optogenetic identification and dopamine neuron stimulation.**

**(A)** Optogenetic identification of D1- and D2-SPNs using “optopatch method” in D2-Cre x Ai32 mice. Increasing light intensities of 488 nm light (0.5, 1.0, 1.5 mW) delivered through the patch pipette produced no response in putative D1-SPNs (top) and progressive membrane depolarization and action potential firing in an example D2-SPN (bottom).

**(B)** Intrinsic properties of D1- and D2-SPNs recorded in this configuration. Left: D2-SPNs show a similar resting V_m_ as D1 SPNs (T-test, *P* = 0.58). Middle: D2-SPNs showed a higher membrane resistance than D1-SPNs (Mann-Whitney test, *P* = 0.008). Right: D2-SPNs had a significantly lower rheobase than D1-SPNs (T-test, *P* < 0.0001). D1: n = 32 cells from 17 mice; D2: n = 29 cells from 17 mice.

**(C)** Schematic of the *ex vivo* patch-clamp recording configuration to confirm optogenetic stimulation of SNc. Whole-cell recordings were performed from ChRmine-expressing SNc dopamine neurons in acute brain slices during 595 nm LED illumination.

**(D)** Representative current-clamp traces showing action potential firing in ChRmine-expressing SNc dopamine neurons in response to light stimulation at 10 Hz / 500 ms (left), 20 Hz / 500 ms (middle), and 20 Hz / 1 s (right), confirming reliable and frequency-matched neuronal firing for 500ms but not 1 s.

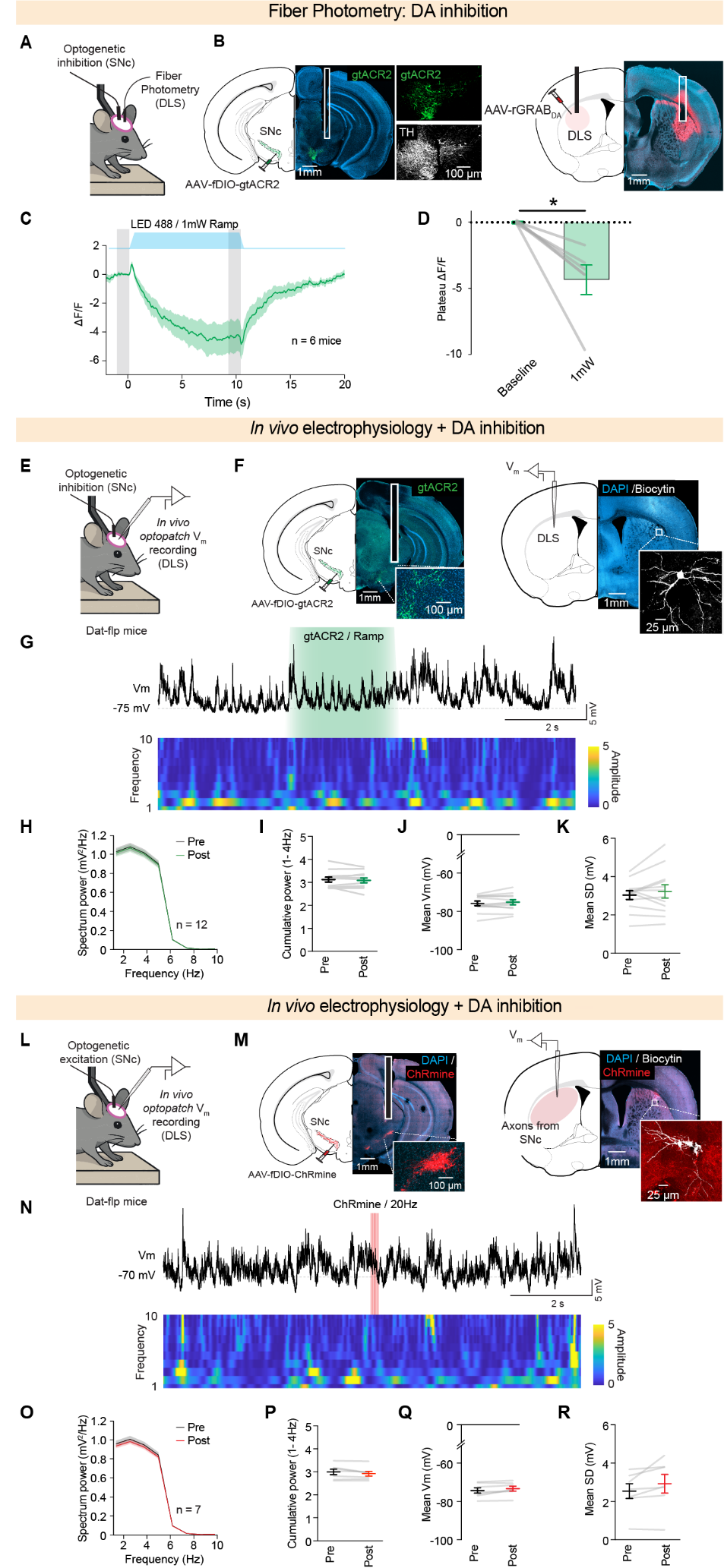

**Figure S2. Bidirectional manipulation of dopamine produces little change in V_m_ dynamics *in vivo.***

**(A)** Schematic of the fiber photometry configuration for dopamine inhibition validation. AAV-fDIO-gtACR2 was expressed in SNc dopamine neurons of DAT-Flp mice and AAV-rGRAB-DA was expressed in the DLS with a fiber optic cannula placed over the ipsilateral DLS.

**(B)** Representative histology showing viral expression of AAV-fDIO-gtACR2-eYFP in SNc dopamine neurons with TH immunostaining confirming expression in dopaminergic neurons, and AAV-rGRAB-DA expression in the DLS with fiber optic placement.

**(C)** Average fiber photometry trace of dopamine release in the DLS during a 10-second ramp light protocol (blue shaded area, LED 488 nm, 1 mW). Dopamine inhibition is sustained throughout the stimulation period and returns to baseline after light offset. n = 6 mice. The grey shaded area indicates the region over which dopamine signals were quantified in (D). Shaded area represents SEM.

**(D)** Plateau ΔF/F during baseline and 1 mW ramp stimulation, confirming reliable and sustained dopamine suppression (Wilcoxon-paired test, *P* = 0.03; n = 6 mice).

**(E)** Schematic of recording configuration. AAV-fDIO-gtACR2 was expressed in SNc dopamine neurons of DAT-Flp mice and whole-cell patch-clamp recordings were performed in the DLS of head-fixed awake mice.

**(F)** Representative histology showing viral expression of AAV-fDIO-gtACR2 in dopaminergic neurons of the SNc and example of a biocytin-filled neuron recorded in the DLS aligned to Paxinos mouse brain atlas.

**(G)** Representative *in vivo* V_m_ trace (top) and time-frequency analysis (bottom) during optogenetic inhibition of SNc dopamine neurons (green shaded area, gtACR2 ramp protocol shown in C).

**(H)** Power spectrum density of V_m_ recordings before and after dopamine inhibition (n = 12 cells from 7 mice).

**(I)** Cumulative power in the 1-4 Hz band before and after dopamine inhibition (T-test, *P* = 0.45).

**(J)** Mean V_m_ before and after dopamine inhibition (T-test, *P* = 0.74).

**(K)** Standard deviation of V_m_ before and after dopamine inhibition (T-test, *P* = 0.65).

**(L)** Schematic of recording configuration. AAV-fDIO-ChRmine was expressed in SNc dopamine neurons of DAT-Flp mice and whole-cell patch-clamp recordings were performed in the DLS of head-fixed mice.

**(M)** Representative histology showing viral expression of AAV-fDIO-ChRmine-mScarlet in dopaminergic neurons of the SNc and ChRmine-expressing SNc axons in the DLS with a biocytin-filled recorded neuron.

**(N)** Representative *in vivo* V_m_ trace (top) and time-frequency analysis (bottom) before, during, and after optogenetic activation of SNc dopamine neurons at 20 Hz (red shaded area).

**(O)** Power spectral density of V_m_ recordings before and after dopamine activation (n = 7 cells from 4 mice).

**(P)** Cumulative power in the 1-4 Hz band before and after dopamine activation (Mann-Whitney, *P* = 0.62).

**(Q)** Mean V_m_ before and after dopamine activation (Mann-Whitney, *P* = 0.53).

**(R)** Standard deviation of V_m_ before and after dopamine activation (Mann-Whitney, *P* = 0.38).

Data are shown as mean ± SEM. Each line circle represents an individual neuron.

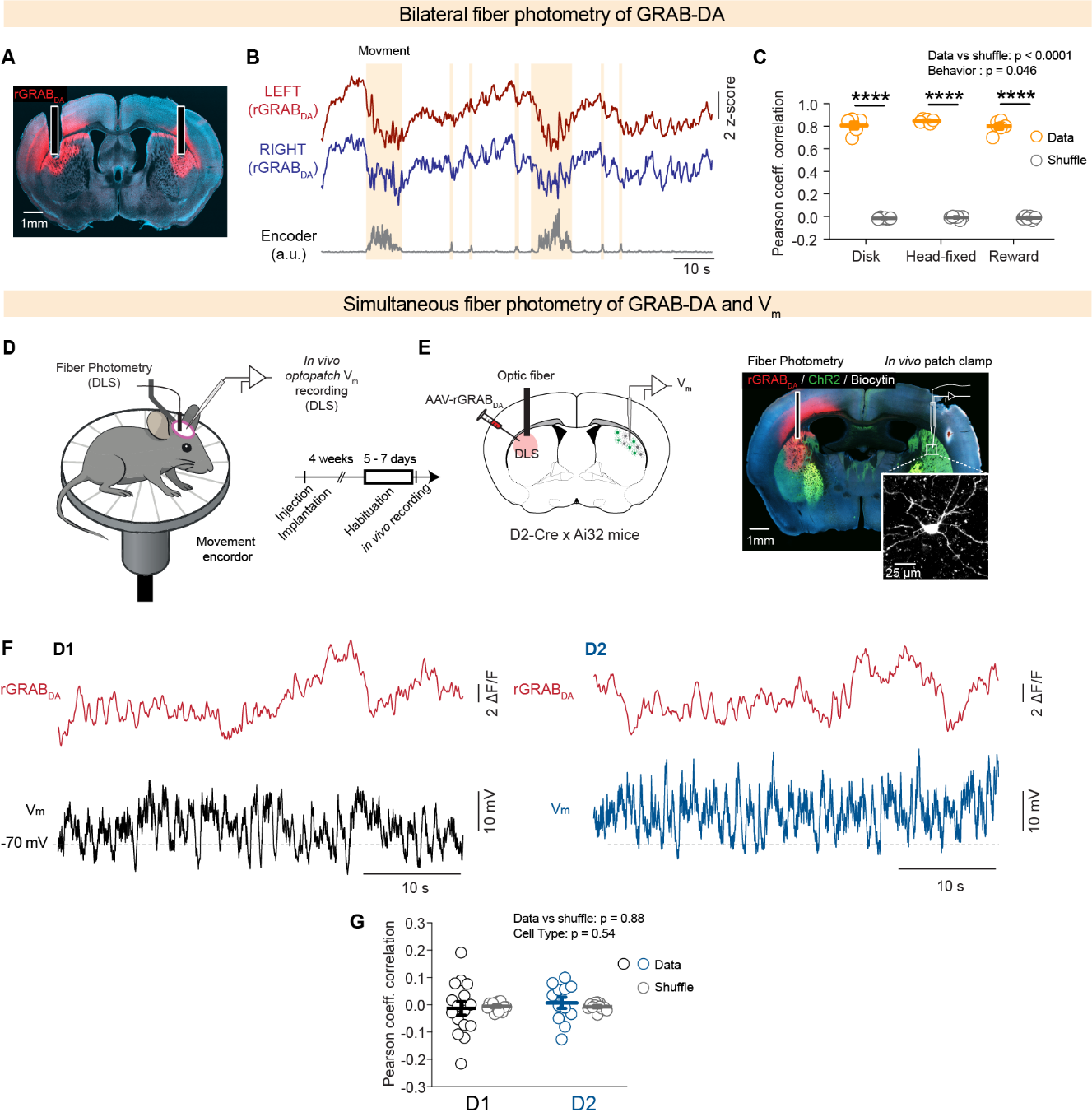

**Fig. S3. Dopamine dynamics are not correlated with SPN V_m_ dynamics.**

**(A)** Representative histology showing bilateral rGRAB-DA expression in the DLS with fiber optic cannulas placed over both hemispheres.

**(B)** Representative simultaneous fiber photometry traces of dopamine release in the left (red) and right (blue) DLS during locomotion bouts (orange shaded areas), with movement encoder signal shown below.

**(C)** Pearson correlation coefficient between left and right hemisphere dopamine signals across three behavioral conditions: disk locomotion, head-fixed platform, and reward delivery, compared to shuffled controls. Dopamine signals are highly correlated between hemispheres in all conditions (two-way ANOVA, data vs shuffle: *P* < 0.0001, behavior: *P* = 0.046; n = 6 mice).

**(D)** Schematic of the recording configuration. Fiber photometry was performed in the DLS while simultaneous V_m_ recordings were performed in the contralateral DLS of head-fixed mice placed on a disk equipped with a movement encoder. Experimental timeline showing injection, implantation, habituation and recording phases.

**(E)** Schematic showing AAV-rGRAB-DA expression and fiber optic placement in the DLS of D2-Cre x Ai32 mice (left). Representative histology showing rGRAB-DA expression, ChR2 expression in D2 neurons, and a biocytin-filled neuron recorded *in vivo*.

**(F)** Representative simultaneous rGRAB-DA fiber photometry traces (top, red) and *in vivo* V_m_ recordings (bottom) during quiet periods in D1-SPNs (left, black) and D2-SPNs (right, blue).

**(J)** Pearson correlation coefficient between dopamine release and Vm dynamics for data and shuffled controls in D1- and D2-SPNs. No significant correlation was detected in either cell type (two-way ANOVA, data vs shuffle: *P* = 0.88, cell type: *P* = 0.94; D1: n = 16 cells from 8 mice; D2: n = 12 cells from 10 mice).

Data are shown as mean ± SEM. Each open circle represents an individual neuron (electrophysiology) or mouse (fiber photometry). *****P* < 0.0001

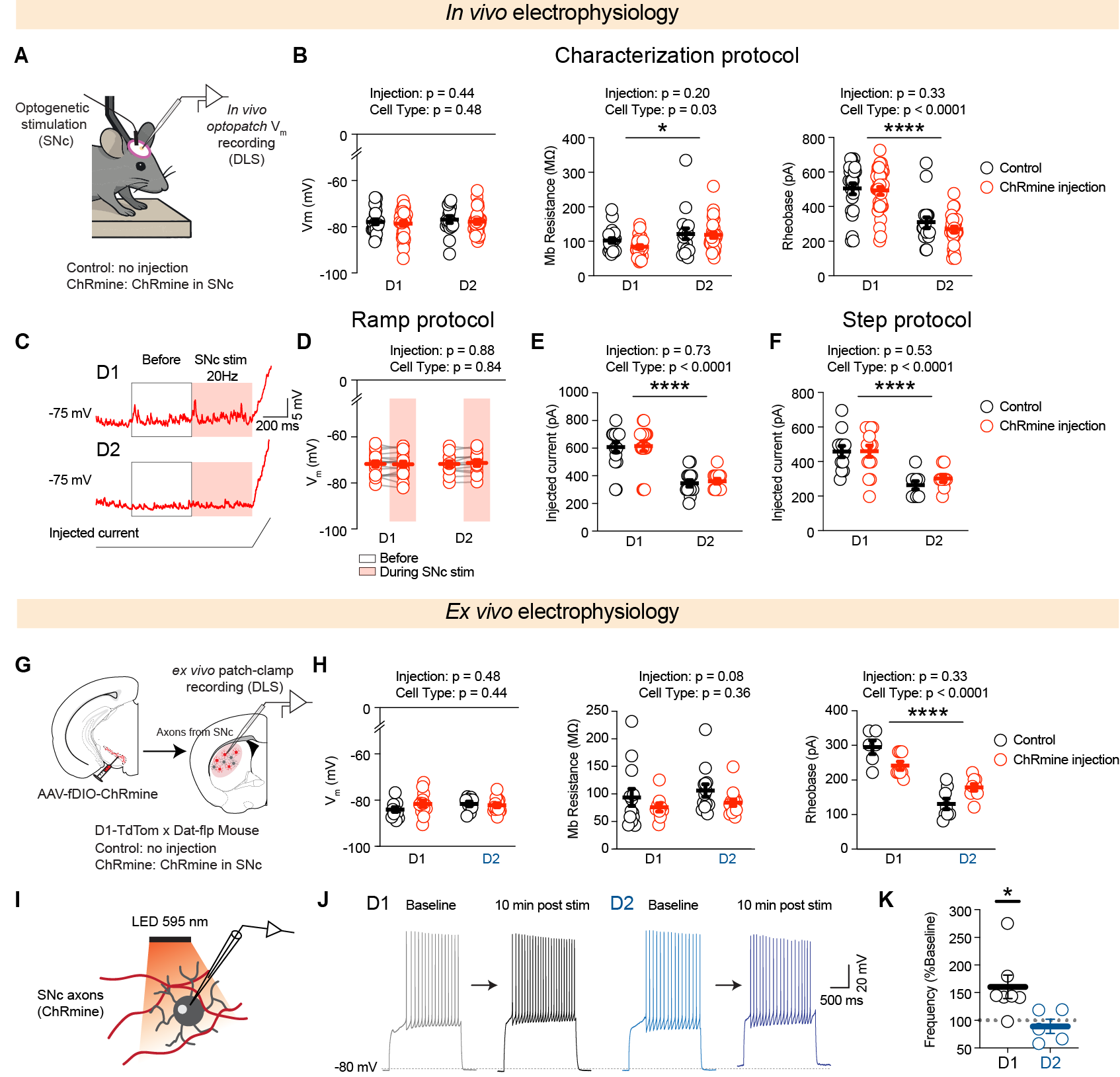

**Fig. S4. ChRmine expression does not alter intrinsic physiology, and dopamine produces persistent increases in D1-SPN excitability ex vivo.**

**(A)** Schematic of the *in vivo* recording configuration. ChRmine was expressed in SNc dopamine neurons of D2-Cre x Ai32 x DAT-Flp mice and V_m_ recordings were performed in the DLS. Control mice received no viral injection in the SNc.

**(B)** Intrinsic properties of D1 and D2-SPNs in control and ChRmine-injected mice. ChRmine-injected cells are the same as Fig.S1. Left: resting membrane potential comparable (two-way ANOVA: injection *P* = 0.44, cell type *P* = 0.48). Middle: membrane resistance was comparable (two-way ANOVA: injection *P* = 0.20, cell type *P* = 0.03). Right: rheobase was comparable between injection conditions (wo-way ANOVA: injection *P* = 0.33, cell type *P* < 0.0001). D1: Control n = 23 cells from 10 mice; ChRmine n = 32 cells from 17 mice; D2: Control n = 17 cells from 9 mice; ChRmine n = 29 cells from 17 mice.

**(C)** Representative *in vivo* current ramp traces in D1-SPNs (top) and D2-SPNs (bottom) before and during SNc stimulation at 20 Hz during the ramp protocol.

**(D)** Resting V_m_ before and during SNc stimulation in D1- and D2-SPNs for control and ChRmine mice (two-way ANOVA; injection *P* = 0.20, cell type *P* = 0.03; D1: n = 19 cells from 14 mice; D2: n = 13 cells from 11 mice).

**(E)** Current injected for the ramp protocol in D1 and D2-SPNs for control and ChRmine mice (two-way ANOVA; injection *P* = 0.73, cell type *P* < 0.0001; D1: Control n = 17 cells from 10 mice; ChRmine: n = 19 cells from 14 mice; D2: Control n = 14 cells from 8 mice; ChRmine: n = 13 cells from 11 mice).

**(F)** Current injected during depolarization protocol (see methods) in D1 and D2-SPNs for control and ChRmine mice (two-way ANOVA; injection *P* = 0.53, cell type *P* < 0.0001; D1: Control n = 12 cells from 6 mice; ChRmine: n = 15 cells from 12 mice; D2: Control n = 9 cells from 7 mice; ChRmine: n = 9 cells from 7 mice).

**(G)** Schematic of the *ex vivo* recording configuration. AAV-fDIO-ChRmine was expressed in SNc dopamine neurons of D1-TdTom x DAT-Flp mice and patch-clamp recordings were performed in DLS slices. Control mice received no viral injection in the SNc.

**(H)** Intrinsic properties of D1 and D2-SPNs in control and ChRmine-injected mice in slice. Left: Resting V_m_ was comparable between conditions (two-way ANOVA: injection *P* = 0.48, cell type *P* = 0.44), Middle: membrane resistance was comparable between conditions (two-way ANOVA: injection *P* = 0.06, cell type *P* = 0.36). Right: rheobase was comparable between conditions (two-way ANOVA: injection *P* = 0.33, cell type). D1: Control n = 14 cells from 4 mice; ChRmine: n = 11 cells from 6 mice; D2: Control n = 14 cells from 4 mice; ChRmine: n = 19 cells from 5 mice.

**(I)** Schematic of the ex vivo stimulation configuration. 595 nm light was delivered onto ChRmine-expressing SNc axons in DLS slices during whole-cell recordings in which current steps were delivered to elicit action potential firing.

**(J)** Representative *ex vivo* current step traces in D1-SPNs (left) and D2-SPNs (right) at baseline and 10 minutes after SNc axon stimulation, showing a persistent increase in action potential frequency in D1-SPNs but not D2-SPNs.

**(K)** Action potential frequency 10 minutes after SNc axon stimulation expressed as percentage of baseline in D- and D2-SPNs (One sample Wilcoxon-test, D1: *P* = 0.03; n = 7 cells from 4 mice; D2: *P* = 0.45; n = 5 cells from 3 mice).

Data are shown as mean ± SEM. Each open circle represents an individual neuron. **P* < 0.05, *****P* < 0.0001.

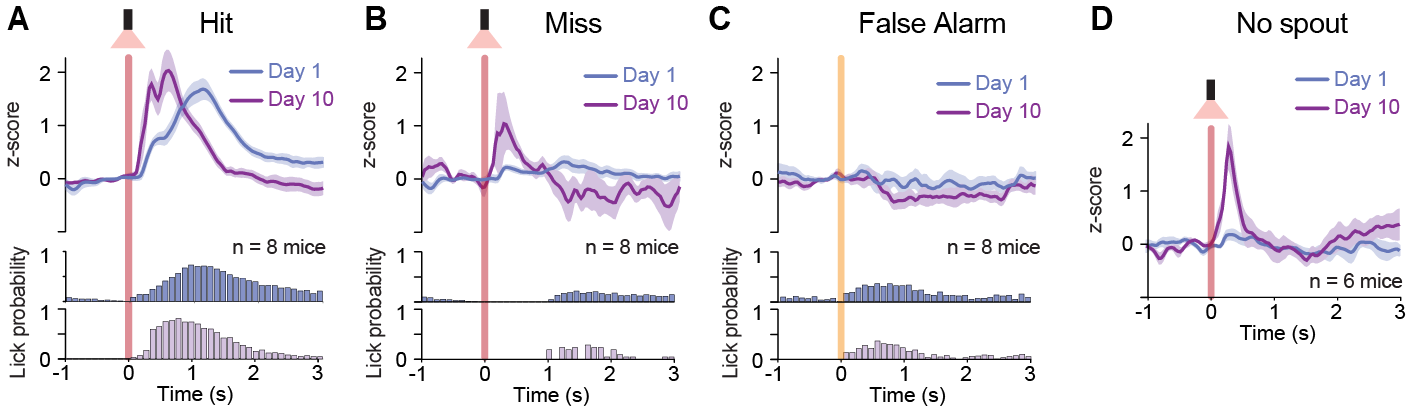

**Fig. S5: Fiber photometry of DA across different trial types in S1 guided learning task**

(A) Grand average of dopamine release aligned S1 stim across hit trials in naïve (day 1) and expert (day 10) mice. Lick probability is shown below each trace. n = 8 mice.
(B) Same as (A), for miss trials.
(C) Same as (A), for false alarm trials, aligned to trial start.
(D) Same as (A), for trials in which the spout was removed (“no spout”). n = 6 mice.

Shaded area represents ± SEM.

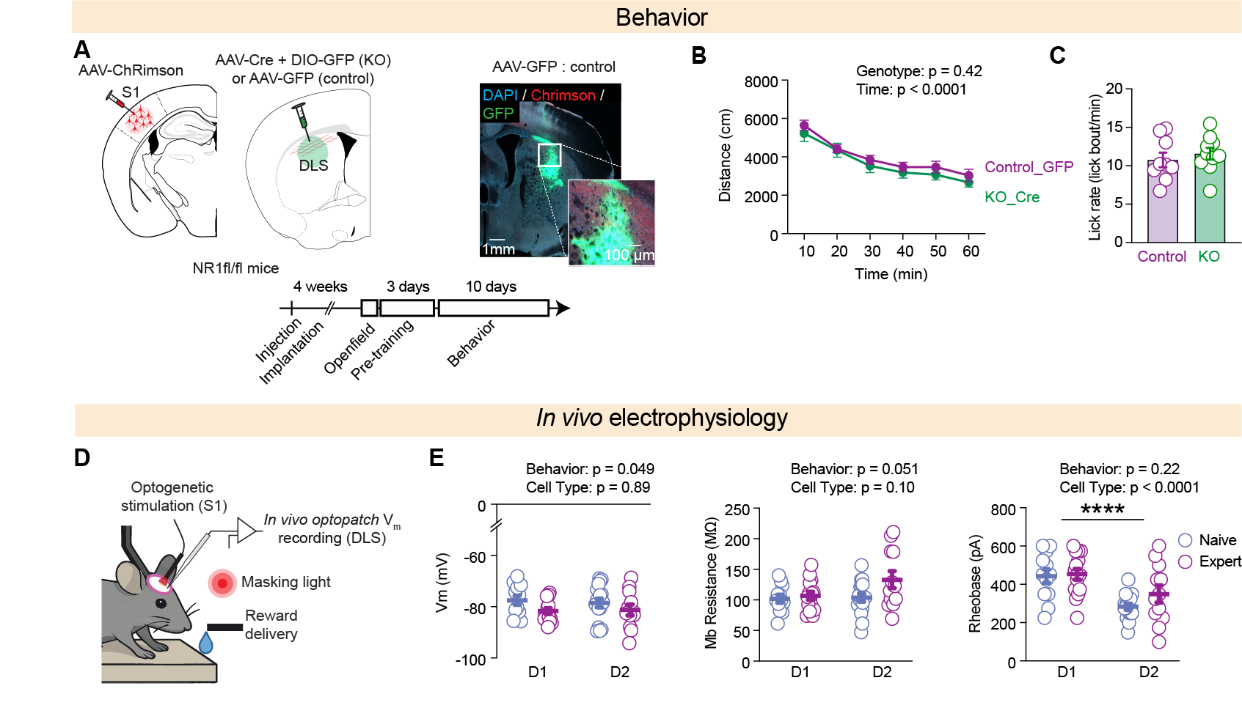

**Fig. S6. Learning does not alter intrinsic membrane properties of D1- or D2-SPNs.**

**(A)** Schematic of the NR1 knockout configuration. AAV-Cre and AAV-DIO-GFP were injected in the DLS of NR1KO-Cre mice to drive NMDA receptor deletion. As a control, NR1KO-Cre mive were injected with AAV-GFP alone. Histology shows GFP expression in the DLS. AAV-ChRimson was expressed in S1 in both groups. Experimental timeline showing injection, open field recording, pre-training and behavioral training phases.

**(B)** Distance traveled over time in the open field in GFP control (purple) and NR1KO conditions (green), confirming that striatal NMDA receptor deletion does not impair locomotion (repeated measure two-way ANOVA: Genotype *P* = 0.42, time *P* < 0.0001; GFP n = 10 mice; KO n = 10 mice).

**(C)** Baseline licking rate during pre-training in control and NR1KO mice, showing no significant difference between groups (Mann-Whitney test, *P* = 0.32; Control: n = 9 mice; KO: n = 10 mice).

**(D)** Schematic of the recording configuration. AAV-ChRimson was expressed in S1 and V_m_ recordings were performed in the DLS of head-fixed awake mice during the behavioral task with reward delivery.

**(E)** Intrinsic properties of D1- and D2-SPNs in naïve (blue) and expert (purple) mice. Left: Resting V_m_ was comparable before and after learning (two-way ANOVA: behavior *P* = 0.049, cell type *P* = 0.051). Middle: membrane resistance was comparable before and after learning (two-way ANOVA: behavior *P* = 0.22, cell type *P* < 0.0001). Right: rheobase was comparable before and after learning (two-way ANOVA: behavior *P* = 0.10, cell type *P* = 0.051) D1: naïve n = 14 cells from 10 mice; expert n = 11 cells from 9 mice; D2: naïve n = 16 cells from 11 mice; expert n = 9 cells from 6 mice.

Data are shown as mean ± SEM. Each open circle represents an individual mouse in C. Each open circle represents an individual neuron in E. *****P* < 0.0001.

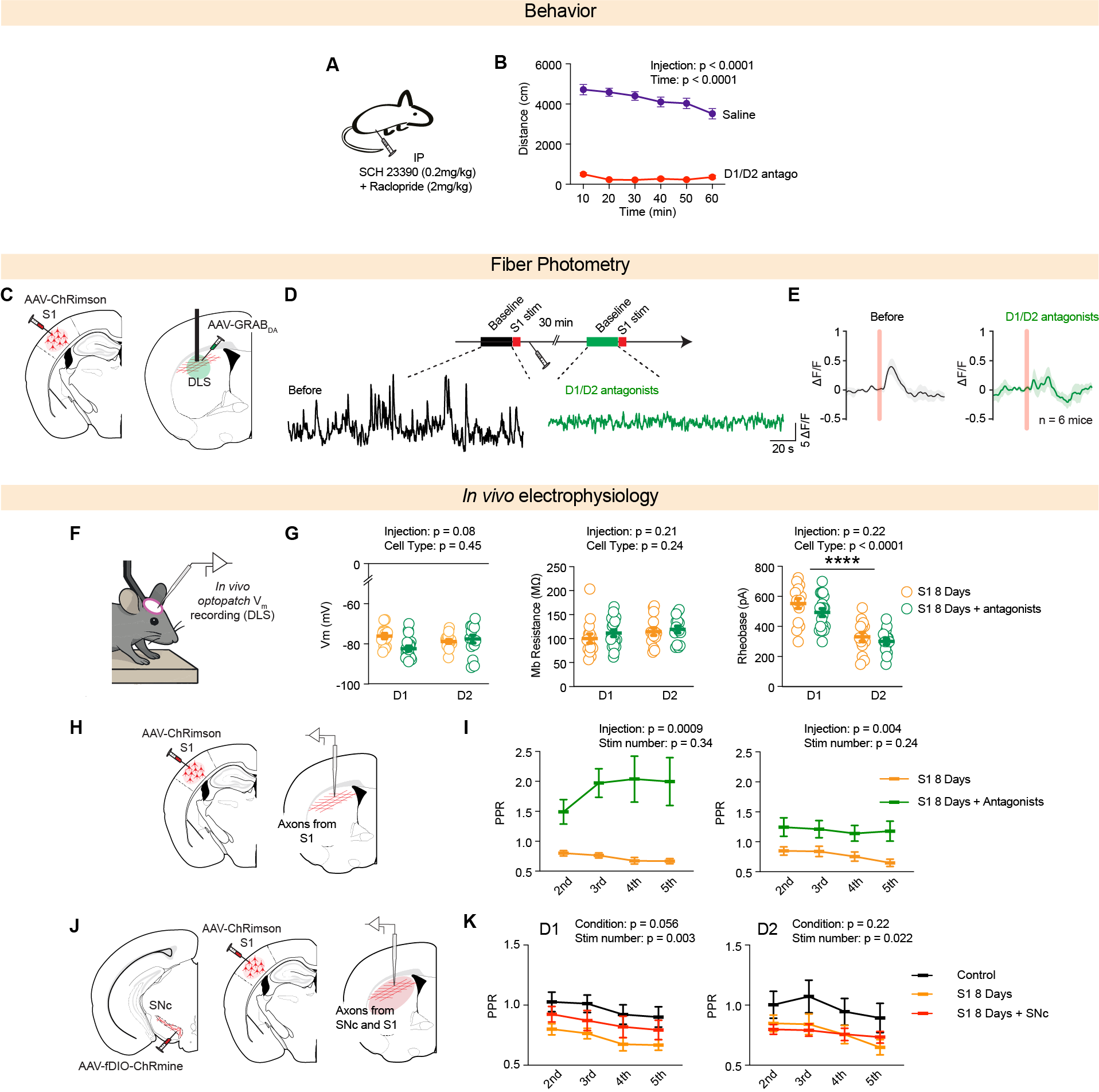

**Fig. S7. Characterization of dopamine modulation on behavior, dopamine release, and corticostriatal synaptic properties**

**(A)** Schematic of the pharmacological approach. Saline or a mix of SCH 23390 (0.2 mg/kg) and raclopride (2 mg/kg) were administered intraperitoneally.

**(B)** Distance traveled over time in the open field following saline or D1/D2 antagonist injection confirming effective dopamine receptor blockade. (repeated measure two-way ANOVA: injection *P* < 0.0001, time *P* < 0.0001; Saline n = 27 mice; antagonist n = 17 mice).

**(C)** Schematic of the fiber photometry configuration for antagonist validation. AAV-ChRimson was expressed in S1 and GRAB-DA was expressed in the DLS with a fiber optic to subsequently measure dopamine release.

**(D)** Representative fiber photometry traces of dopamine fluctuations in the DLS in a head-fixed mouse before (black) and 30 min after D1/D2 antagonist injection (green), showing suppression of dopamine. transients.

**(E)** Average fiber photometry traces of dopamine release aligned to S1 stimulation before (left) and after D1/D2 antagonist injection (right). n = 6 mice.

**(F)** Schematic of the *in vivo* recording configuration for intrinsic property assessment after repeated stimulation of S1with or without antagonist.

**(G)** Intrinsic properties of D1- and D2-SPNs in S1 8 days (yellow) and S1 8 days with antagonist (green) conditions. Left: Resting membrane potential was comparable between conditions (two-way ANOVA: injection *P* = 0.06, cell type *P* = 0.45). Middle: membrane resistance was comparable between conditions (two-way ANOVA: *P* = 0.21, cell type *P* = 0.24). Right: rheobase was comparable between conditions, with a significant difference in cell type (two-way ANOVA: injection *P* = 0.22, cell type *P* < 0.0001). D1: S1 8 days n = 15 cells from 5 mice; S1 8 days + antagonist n = 19 cells from 8 mice; D2: S1 8 days n = 15 cells from 5 mice; S1 8 days + antagonist n = 15 cells from 7 mice.

**(H)** Schematic of the *in vivo* recording configuration for paired-pulse ratio assessment. AAV-ChRimson was expressed in S1 and V_m_ recordings were performed in the DLS.

**(I)** Paired-pulse ratio across successive stimulations in D1-SPNs (left) and D2-SPNs (right) in S1 8 days and (yellow) S1 8 days with antagonist conditions (green, two-way ANOVA; D1: injection *P* = 0.0009, stimulation number *P* = 0.34; D2: injection *P* = 0.004, stimulation number *P* = 0.24; S1 8 days n = 14 cells from 5 mice; S1 8 days + antagonist n = 17 cells from 8 mice; D2: S1 8 days n = 13 cells from 5 mice; S1 8 days + antagonist n = 11 cells from 6 mice).

**(J)** Schematic of the recording configuration. AAV-fDIO-ChRmine was expressed in SNc dopamine neurons and AAV-ChRimson was expressed in S1 of D2-Cre x Ai32 x DAT-Flp mice, allowing independent co-stimulation of cortical inputs and dopamine neurons. V_m_ recordings were performed in the DLS.

**(K)** Paired-pulse ratio across successive stimulations in D1-SPNs (left) and D2-SPNs (right) across the three conditions: control (black), S1 8 days stimulation (yellow), and S1 8 days stimulation plus SNc stimulation (red). (two-way ANOVA; D1: condition *P* = 0.056, stimulation number *P* = 0.003; D2: condition *P* = 0.22, stimulation number *P* = 0.022; D1: control: n = 27 cells from 17 mice; S1 8 days n = 14 cells from 5 mice; S1 + SNc 8 days n = 15 cells from 7 mice; S1 8 days + antagonist n = 17 cells from 8 mice; D2: *P* < 0.0001; control: n = 29 cells from 17 mice; S1 8 days n = 13 cells from 5 mice; S1 + SNc 8 days n = 15 cells from 10 mice; S1 8 days + antagonist n = 11 cells from 6 mice).

Data are shown as mean ± SEM. Each open circle represents an individual neuron. *****P* < 0.0001.

**Supplementary Tables**

|  | n | N | Pre | Stim | Post | Test | p value | Significant |
| --- | --- | --- | --- | --- | --- | --- | --- | --- |
| In vivo electrophysiology - ChRmine stimulation | | | | | | | | |
| **Amplitude (pA)** | | |  | | | | | |
| D1 | 9 | 6 | 3.81 ± 1.22 | 3.76 ± 1.17 | 4.83 ± 1.41 | RM One-way ANOVA | 0.006 | * |
| *Pre vs stim* |  | | | | | Sidak's post-hoc test | 0.27 | n.s. |
| *Stim vs Post* |  | | | | | Sidak's post-hoc test | 0.0065 | ** |
| *Pre vs Post* |  | | | | | Sidak's post-hoc test | 0.18 | n.s. |
| D2 | 9 | 6 | 2.64 ± 0.53 | 2.28 ± 0.45 | 2.94 ± 0.84 | RM One-way ANOVA | 0.44 | n.s. |
| **PPR** | | |  | | | | | |
| D1 | 9 | 6 | 3.81 ± 1.22 | 3.76 ± 1.17 | 4.83 1.41 | RM One-way ANOVA | 0.86 | n.s. |
| D2 | 9 | 6 | 1.28 ± 0.51 | 1.42 ± 0.49 | 1.2 ± 0.28 | RM One-way ANOVA | 0.82 | n.s. |
| Fiber Photometry - ChRmine stimulation | | | | | | | | |
| **Peak amplitude (ΔF/F)** | | |  | | | RM One-way ANOVA | 0.038 | * |
| *Reward* | | 6 |  | 10.06 ± 0.94 |  |  | | |
| *2mW 10Hz vs Reward* | | 6 |  | 5.816 ± 0.96 |  | Sidak's post-hoc test | 0.038 | * |
| *2mW 20Hz vs Reward* | | 6 |  | 10.16 ± 1.22 |  | Sidak's post-hoc test | 0.999 | n.s. |
| *4mW 20Hz vs Reward* | | 6 |  | 14.56 ± 1.12 |  | Sidak's post-hoc test | 0.02 | * |

**Table S1. Summary table of *in vivo* whole cell recording and fiber photometry (Fig. 1)**

N refers to the number of mice, n refers to the number of cells. All data are presented as mean ± SEM. N.s., non-significant; * *P* < 0.05.

|  | n | N | D1 | n | N | D2 | Test | p value | Significant |
| --- | --- | --- | --- | --- | --- | --- | --- | --- | --- |
| In vivo electrophysiology | | | | | | | | | |
| Resting potential (mV) | 32 | 17 | -78.62 ± 1.1 | 29 | 17 | -77.8 ± 1.0 | T-test | 0.58 | n.s. |
| Membrane resistance (MΩ) | 32 | 17 | 84.9 ± 5 | 29 | 17 | 119.2 ± 8 | Mann-Whitney | 0.008 | *** |
| Rheobase (pA) | 32 | 17 | 493 ± 23 | 29 | 17 | 269.8 ± 17 | T-test | < 0.0001 | **** |

**Table S2. Summary table of *in vivo* whole cell recording (Fig. S1)**

N refers to the number of mice, n refers to the number of cells. All data are presented as mean ± SEM. N.s., non-significant; *** *P* < 0.001, **** *P* < 0.0001.

|  | n | N | Pre | Stim | Post | Test | p value | Significant |
| --- | --- | --- | --- | --- | --- | --- | --- | --- |
| In vivo electrophysiology | | | | | | | | |
| **Rheobase D1 (pA)** | | |  | | | | | |
| Control | 17 | 10 | 486.5 ± 28.2 | 467.2 ± 30.1 | 453.4 ± 30.0 | RM Two-way ANOVA | stim: 0.037 | * |
| ChRmine | 19 | 14 | 499.8 ± 33.8 | 492.9 ± 31.8 | 479.1 ± 28.6 |  | condition: 0.61 | n.s. |
| *Control: Pre vs stim* | | |  | | | *Sidak's post-hoc test* | 0.54 | n.s. |
| *Control: Stim vs Post* | | |  | | | *Sidak's post-hoc test* | 0.82 | n.s. |
| *Control: Pre vs Post* | | |  | | | *Sidak's post-hoc test* | 0.06 | n.s. |
| *ChRmine: Pre vs stim* | | |  | | | *Sidak's post-hoc test* | 0.93 | n.s. |
| *ChRmine: Stim vs Post* | | |  | | | *Sidak's post-hoc test* | 0.75 | n.s. |
| *ChRmine: Pre vs Post* | | |  | | | *Sidak's post-hoc test* | 0.21 | n.s. |
| **Rheobase D2 (pA)** | | |  | | | | | |
| Control | 15 | 8 | 271.8 ± 21.4 | 262.8 ± 17.9 | 263.5 ± 22.7 | RM Two-way ANOVA | stim: 0.64 | n.s. |
| ChRmine | 13 | 11 | 266.5 ± 19.8 | 267.6 ± 19.7 | 263.5 ± 18.5 |  | condition : 0.99 | n.s. |
| **Frequency D1 (Hz)** | | |  | | | | | |
| Control | 12 | 7 | 28.67 ± 2.4 | 30.25 ± 3.4 | 31.85 ± 2.7 | RM Two-way ANOVA | stim: 0.037 | * |
| ChRmine | 15 | 12 | 23.75 ± 2.6 | 23.83 ± 3.1 | 24.27 ± 2.7 |  | condition: 0.61 | n.s. |
| **Frequency D2 (Hz)** | | |  | | | | | |
| Control | 9 | 7 | 25.89 ± 2.8 | 27.56 ± 2.7 | 24.14 ± 2.7 | RM Two-way ANOVA | stim: 0.64 | n.s. |
| ChRmine | 9 | 7 | 21.36 ± 3.1 | 23.5 ± 3.2 | 23.19 ± 2.5 |  | condition : 0.99 | n.s. |
| Ex vivo electrophysiology | | | | | | | | |
| **Frequency D1 (Hz)** | | |  | | | | | |
| Control | 14 | 4 | 8.70 ± 0.6 | 8.11 ± 0.4 | 9.10 ± 0.6 | RM Two-way ANOVA | stim: < 0.0001 | **** |
| ChRmine | 11 | 6 | 7.57 ± 0.6 | 8.36 ± 0.6 | 10.61 ± 0.9 |  | condition: 0.78 | n.s. |
| *Control: Pre vs stim* | | |  | | | *Sidak's post-hoc test* | 0.53 | n.s. |
| *Control: Stim vs Post* | | |  | | | *Sidak's post-hoc test* | 0.07 | n.s. |
| *Control: Pre vs Post* | | |  | | | *Sidak's post-hoc test* | 0.9 | n.s. |
| *ChRmine: Pre vs stim* | | |  | | | *Sidak's post-hoc test* | 0.32 | n.s. |
| *ChRmine: Stim vs Post* | | |  | | | *Sidak's post-hoc test* | < 0.0001 | **** |
| *ChRmine: Pre vs Post* | | |  | | | *Sidak's post-hoc test* | < 0.0001 | **** |
| **Frequency D2 (Hz)** | | |  | | | | | |
| Control | 14 | 4 | 8.21 ± 0.6 | 7.57 ± 0.4 | 8.32 ± 0.7 | RM Two-way ANOVA | stim: 0.64 | n.s. |
| ChRmine | 19 | 5 | 8.72 ± 0.7 | 8.53 ± 0.9 | 8.74 ± 0.9 |  | condition : 0.99 | n.s. |

**Table S3. Summary table of *in vivo* and *ex vivo* whole cell recording (Fig. 2)**

N refers to the number of mice, n refers to the number of cells. All data are presented as mean ± SEM. n.s. non significant; * *P* < 0.05; **** *P* < 0.0001.

|  | n | N | Before | After | Test | p value | Significant |
| --- | --- | --- | --- | --- | --- | --- | --- |
| Fiber photometry | | | | | | | |
| Ramp inhibition |  | 6 | 0.008 ± 0.05 | - 4.31 ± 1.12 | Wilcoxon-paired test | 0.011 | * |
| (ΔF/F) |  |  |  |  |  |  |  |
| In vivo electrophysiology - Inhibition (gtACR2) | | | | | | | |
| Cumulative power | 12 | 7 | 3.12 ± 0.1 | 3.09 ± 0.1 | T-test | 0.82 | n.s. |
| (1-4Hz) |  |  |  |  |  |  |  |
| Mean Vm (mV) | 12 | 7 | -75.88 ± 1.2 | -75.27 ± 1.3 | T-test | 0.74 | n.s. |
| Mean SD (mV) | 12 | 7 | 1.42 ± 0.23 | 3.23 ± 0.35 | T-test | 0.65 | n.s. |
| In vivo electrophysiology - Excitation (ChRmine) | | | | | | | |
| Cumulative power | 7 | 4 | 3.0 ± 0.12 | 2.93 ± 0.11 | Mann-Whitney | 0.62 | n.s. |
| (1-4Hz) |  |  |  |  |  |  |  |
| Mean Vm (mV) | 7 | 4 | -74.18 ± 1.38 | -73.2 ± 1.3 | Mann-Whitney | 0.53 | n.s. |
| Mean SD (mV) | 7 | 4 | 2.52 ± 0.39 | 2.91 ± 0.49 | Mann-Whitney | 0.38 | n.s. |

**Table S4. Summary table of *in vivo* whole cell recording and fiber photometry (Fig. S2)**

N refers to the number of mice, n refers to the number of cells. All data are presented as mean ± SEM. N.s., non-significant; * *P* < 0.05.

|  | n | N | Data | n | N | shuffle | Test | p value | Significant |
| --- | --- | --- | --- | --- | --- | --- | --- | --- | --- |
| Bilateral fiber photometry | | | | | | | | | |
| Pearson coefficient Correlation (r) | | |  | | | | Two-way RM ANOVA | Behavior type: 0.046 | * |
|  |  |  |  |  |  |  |  | Data vs shuffle < 0.0001 | **** |
| Disk | | 6 | 0.81 ± 0.03 |  |  | -0.014 ± 0.002 | Sidak's post-hoc test | < 0.0001 | **** |
| Head-fixed | | 6 | 0.85 ± 0.005 |  |  | -0.007 ± 0.005 | Sidak's post-hoc test | < 0.0001 | **** |
| Reward | | 6 | 0.80 ± 0.022 |  |  | -0.012 ± 0.006 | Sidak's post-hoc test | < 0.0001 | **** |
| Simultaneous *in vivo* electrophysiology and fiber photometry | | | | | | | | | |
| Pearson coefficient Correlation (r) | | |  | | | | Two-way RM ANOVA | Cell type : 0.94 | n.s. |
|  |  |  |  |  |  |  |  | Data vs shuffle: 0.88 | n.s. |
| *Data* | 16 | 8 | -0.013 ± 0.025 | 12 | 10 | 0.008 ± 0.02 |  | | |
| *Shuffle* | 16 | 8 | -0.005 ± 0.004 | 12 | 10 | -0.007 ± 0.004 |  | | |

**Table S5. Summary table of *in vivo* whole cell recording and fiber photometry (Fig. S3)**

N refers to the number of mice, n refers to the number of cells. All data are presented as mean ± SEM. N.s., non-significant; * *P* < 0.05, **** *P* < 0.0001.

|  | n | N | D1 | n | N | D2 | Test | p value | Significant |
| --- | --- | --- | --- | --- | --- | --- | --- | --- | --- |
| In vivo electrophysiology | | | | | | | | | |
| Resting potential (mV) | | |  | | | | RM Two-way ANOVA | injection p = 0.48 | n.s. |
|  |  |  |  |  |  |  |  | cell type: p = 0.44 | n.s. |
| Control | 23 | 10 | -77.89 ± 1.1 | 17 | 9 | -76.87 ± 1.5 |  | | |
| ChRmine | 32 | 17 | -78.62 ± 1.1 | 29 | 17 | -77.8 ± 1.0 |  | | |
| Membrane resistance (MΩ) | | |  | | | | RM Two-way ANOVA | injection p = 0.20 | n.s. |
|  |  |  |  |  |  |  |  | cell type: p = 0.0026 | ** |
| Control | 23 | 10 | 103.7 ± 6.8 | 17 | 9 | 122.8 ± 16.1 | Sidak's post-hoc test | 0.16 | n.s. |
| ChRmine | 32 | 17 | 84.9 ± 5.0 | 29 | 17 | 119.2 ± 8.1 | Sidak's post-hoc test | 0.002 | ** |
| Rheobase (pA) | | |  | | | | RM Two-way ANOVA | injection p = 0.33 | n.s. |
|  |  |  |  |  |  |  |  | cell type: p < 0.0001 | **** |
| Control | 23 | 10 | 504.3 ± 31.4 | 17 | 9 | 308.8 ± 31.7 | Sidak's post-hoc test | < 0.0001 | **** |
| ChRmine | 32 | 17 | 493 ± 22.7 | 29 | 17 | 269.8 ± 17.0 | Sidak's post-hoc test | < 0.0001 | **** |
| Ramp protocol: Vm before / during light | | |  | | | | RM Two-way ANOVA | light stim: p = 0.88 | n.s. |
|  |  |  |  |  |  |  |  | cell type: p = 0.84 | n.s. |
| Before | 19 | 14 | -71.8 ± 1.1 | 13 | 11 | -71.8 ± 1.2 |  | | |
| During | 19 | 14 | -71.9 ± 1.2 | 13 | 11 | -71.4 ± 1.3 |  | | |
| Ramp protocol: Injected current (pA) | | |  | | | | RM Two-way ANOVA | injection p = 0.73 | n.s. |
|  |  |  |  |  |  |  |  | cell type: p < 0.0001 | **** |
| Control | 17 | 10 | 608.8 ± 34.4 | 14 | 8 | 346.4 ± 23.7 | Sidak's post-hoc test | < 0.0001 | **** |
| ChRmine | 19 | 14 | 615.8 ± 35.3 | 13 | 11 | 361.5 ± 18.0 | Sidak's post-hoc test | < 0.0001 | **** |
| Step protocol: Injected current (pA) | | |  | | | | RM Two-way ANOVA | injection p = 0.53 | n.s. |
|  |  |  |  |  |  |  |  | cell type: p < 0.0001 | **** |
| Control | 12 | 6 | 462.5 ± 32.6 | 9 | 7 | 266.7 ± 23.6 | Sidak's post-hoc test | 0.0002 | *** |
| ChRmine | 15 | 12 | 463.3 ± 32.5 | 9 | 7 | 305.6 ± 21.2 | Sidak's post-hoc test | 0.0015 | ** |
| ex vivo electrophysiology | | | | | | | | | |
| Resting potential (mV) | | |  | | | | RM Two-way ANOVA | injection p = 0.48 | n.s. |
|  |  |  |  |  |  |  |  | cell type: p = 0.44 | n.s. |
| Control | 12 | 4 | -84.0 ± 1.0 | 9 | 4 | -81.7 ± 1.0 |  | | |
| ChRmine | 11 | 6 | -81.6 ± 1.4 | 14 | 5 | -82.3 ± 0.8 |  | | |
| Resting potential (mV) | | |  | | | | RM Two-way ANOVA | injection p = 0.20 | n.s. |
|  |  |  |  |  |  |  |  | cell type: p = 0.0026 | ** |
| Control | 12 | 4 | 93.6 ± 15.6 | 9 | 4 | 106.3 ± 11.3 | Sidak's post-hoc test | 0.16 | n.s. |
| ChRmine | 11 | 6 | 75.82 ± 7.4 | 14 | 5 | 84.2 ± 5.9 | Sidak's post-hoc test | 0.002 | ** |
| Resting potential (mV) | | |  | | | | RM Two-way ANOVA | injection p = 0.33 | n.s. |
|  |  |  |  |  |  |  |  | cell type: p < 0.0001 | **** |
| Control | 12 | 4 | 293.3 ± 19.1 | 9 | 4 | 130 ± 15.6 | Sidak's post-hoc test | < 0.0001 | **** |
| ChRmine | 11 | 6 | 240 ± 12.0 | 14 | 5 | 177.8 ± 9.7 | Sidak's post-hoc test | < 0.0001 | **** |

**Table S6. Summary table of *in vivo* and *ex vivo* whole cell recording (Fig. S4)**

N refers to the number of mice, n refers to the number of cells. All data are presented as mean ± SEM. n.s. non significant; ** *P* < 0.01; **** *P* < 0.0001.

|  | N | Naïve | Expert | Test | p value | Significant |
| --- | --- | --- | --- | --- | --- | --- |
| Behavior | | | | | | |
| Light overS1 | 12 | Hit: 0.94 ± 0.01 | FA: 0.23 ± 0.03 | RM Two-way ANOVA | fiber loc: < 0.0001 | **** |
| Light shifted | 12 | Hit: 0.36 ± 0.05 | FA: 0.27 ± 0.04 |  | Hit vs FA: < 0.0001 | **** |
| *Hit: S1 vs Shifted* | |  | | | < 0.0001 | **** |
| *FA: S1 vs Shifted* | |  | | | 0.57 | n.s. |
| D' | 16 | 0.26 ± 0.11 | 2.49 ± 0.14 | Paired T-test | < 0.0001 | **** |
| Reaction time (s) | 16 | 0.57 ± 0.03 | 0.36 ± 0.02 | Paired T-test | < 0.0001 | **** |
| Reaction SD (s) | 16 | 0.28 ± 0.01 | 0.11 ± 0.01 | Paired T-test | < 0.0001 | **** |
| Fiber Photometry | | | | | | |
| **Hit trial** | |  | | | | |
| Max Peak (z-score) | 8 | 1.85 ± 0.16 | 2.60 ± 0.38 | Paired T-test | 0.02 | * |
| Slope (z-score/s) | 8 | 5.08 ± 0.51 | 20.46 ± 4.68 | Wilcoxon matched-pairs | 0.008 | ** |
| **No spout** | |  | | | | |
| Max Peak (z-score) | 6 | 0.353 ± 0.08 | 2.14 ± 0.33 | Wilcoxon matched-pairs | 0.31 | * |
| Slope (z-score/s) | 6 | 3.74 ± 0.98 | 21.92 | Wilcoxon matched-pairs | 0.31 | * |

**Table S7. Summary table of behavior and fiber photometry (Fig. 3)**

N refers to the number of mice. All data are presented as mean ± SEM. n.s. non significant; * *P* < 0.05; ** *P* < 0.01; **** *P* < 0.0001.

|  | n | N | Naïve | n | N | Expert | Test | p value | Significant |
| --- | --- | --- | --- | --- | --- | --- | --- | --- | --- |
| In vivo electrophysiology | | | | | | | | | |
| D' |  | 12 | 0.24 ± 0.09 |  | 12 | 2.57 ± 0.17 | Paired T-test | < 0.0001 | **** |
| ΔVm - behavior (mV) |  | | | | | | RM Two-way ANOVA | behavior: p = 0.0009 | *** |
|  |  |  |  |  |  |  |  | cell type: p = 0.89 | n.s. |
| D1 | 14 | 10 | 1.45 ± 0.39 | 11 | 9 | 3.39 ± 0.69 | *Sidak's post-hoc test* | 0.027 | * |
| D2 | 16 | 11 | 1.39 ± 0.36 | 9 | 6 | 3.31 ± 0.84 | *Sidak's post-hoc test* | 0.035 | * |
| ΔVm 10Hz stim (mV) |  | | | | | | RM Two-way ANOVA | behavior: p = 0.0005 | *** |
|  |  |  |  |  |  |  |  | cell type: p = 0.31 | n.s. |
| D1 | 12 | 9 | 1.98 ± 0.66 | 12 | 10 | 4.08 ± 0.59 |  | 0.022 | * |
| D2 | 11 | 7 | 1.31 ± 0.17 | 10 | 8 | 3.56 ± 0.72 |  | 0.022 | * |
| PPR D1 | 12 | 9 |  | 12 | 10 |  | RM Two-way ANOVA | behavior: p = 0.043 | * |
|  |  |  |  |  |  |  |  | stim number: p = 0.81 | n.s. |
| *2nd* |  | | 1.08 ± 0.20 |  | | 0.81 ± 0.09 | *Sidak's post-hoc test* | 0.5981 | n.s. |
| *3rd* |  | | 1.07 ± 0.27 |  | | 0.70 ± 0.04 | *Sidak's post-hoc test* | 0.282 | n.s. |
| *4th* |  | | 0.84 ± 0.10 |  | | 0.81 ± 0. 5 | *Sidak's post-hoc test* | 0.9999 | n.s. |
| *5th* |  | | 0.92 ± 0.17 |  | | 0.73 ± 0.09 | *Sidak's post-hoc test* | 0.8331 | n.s. |
| PPR D2 | 10 | 6 |  | 10 | 8 |  | RM Two-way ANOVA | behavior: p = 0.025 | * |
|  |  |  |  |  |  |  |  | stim number: p = 0.45 | n.s. |
| *2nd* |  | | 1.29 ± 0.18 |  | | 0.89 ± 0.07 | *Sidak's post-hoc test* | 0.1317 | n.s. |
| *3rd* |  | | 1.32 ± 0.17 |  | | 1.02 ± 0.09 | *Sidak's post-hoc test* | 0.3734 | n.s. |
| *4th* |  | | 1.06 ± 0.13 |  | | 0.91 ± 0.12 | *Sidak's post-hoc test* | 0.8896 | n.s. |
| *5th* |  | | 0.99 ± 0.13 |  | | 0.98 ± 0.11 | *Sidak's post-hoc test* | >0.9999 | n.s. |
| Ex vivo electrophysiology | | | | | | | | | |
| AMPA / NMDA ratio |  | | | | | | RM Two-way ANOVA | behavior: p < 0.0001 | **** |
|  |  |  |  |  |  |  |  | cell type: p = 0.79 | n.s. |
| D1 | 18 | 8 | 4.79 ± 0.55 | 13 | 7 | 2.68 ± 0.19 |  | 0.022 | ** |
| D2 | 17 | 7 | 4.76 ± 0.43 | 14 | 6 | 2.94 ± 0.27 |  | 0.022 | ** |
|  |  | N | Control |  | N | KO | Test | p value | Significant |
| Behavior NR1KO | | | | | | | | | |
| D' |  | 9 | 2.43 ± 0.41 |  | 10 | 1.06 ± 0.28 | T-test | 0.012 | * |
| Reaction SD (s) |  | 9 | 0.13 ± 0.03 |  | 10 | 0.21 ± 0.03 | T-test | 0.045 | * |

**Table S8.Summary table of *in vivo* and *ex vivo* whole cell recording and behavior in NR1KO (Fig. 4)**

N refers to the number of mice, n refers to the number of cells. All data are presented as mean ± SEM. N.s., non-significant; * *P* < 0.05, ** *P* < 0.01; *** *P* < 0.001; **** *P* < 0.0001.

|  | | N | Control | | N | KO | Test | p value | Significant |
| --- | --- | --- | --- | --- | --- | --- | --- | --- | --- |
| Behavior | | | | | | | | | |
| Locomotion | | 9 |  | | 10 |  | RM Two-way ANOVA | injection p = 0.42 | n.s. |
|  |  |  |  |  |  |  |  | time: p < 0.0001 | **** |
| Lick rate | | 9 | 10.8 ± 0.9 | | 10 | 11.6 ± 0.8 | Mann-Whitney | 0.36 | n.s. |
|  | n | N | D1 | n | N | D2 | Test | p value | Significant |
| In vivo electrophysiology | | | | | | | | | |
| Resting potential (mV) | | |  | | | | RM Two-way ANOVA | behavior p = 0.048 | * |
|  |  |  |  |  |  |  |  | cell type: p = 0.88 | n.s. |
| Naïve | 14 | 10 | -77.5 ± 1.8 | 16 | 11 | -78.45 ± 1.7 |  | | |
| Expert | 11 | 9 | -81.67 ± 1.2 | 9 | 6 | -81.22 ± 2.1 |  | | |
| *D1: Naïve vs expeprt* | | |  | | | | *Sidak's post-hoc test* | 0.18 | n.s. |
| *D2: Naïve vs expeprt* | | |  | | | | *Sidak's post-hoc test* | 0.44 | n.s. |
| Membrane resistance (MΩ) | | |  | | | | RM Two-way ANOVA | behavior p = 0.051 | n.s. |
|  |  |  |  |  |  |  |  | cell type: p = 0.098 | n.s. |
| Naïve | 14 | 10 | 102 ± 6.5 | 16 | 11 | 104 ± 6.6 |  | | |
| Expert | 11 | 9 | 106.7 ± 6.4 | 9 | 6 | 133.3 ± 13.8 |  | | |
| Rheobase (pA) | | |  | | | | RM Two-way ANOVA | behavior p = 0.22 | n.s. |
|  |  |  |  |  |  |  |  | cell type: p < 0.0001 | **** |
| Naïve | 14 | 10 | 443.8 ± 35.9 | 16 | 11 | 283.8 ± 15.5 | Sidak's post-hoc test | 0.0009 | *** |
| Expert | 11 | 9 | 453.3 ± 28.1 | 9 | 6 | 350 ± 44.5 | Sidak's post-hoc test | 0.043 | * |

**Table S9. Summary table of behavior and *in vivo* whole cell recording (Fig. S6)**

N refers to the number of mice, n refers to the number of cells. All data are presented as mean ± SEM. N.s., non-significant; * *P* < 0.05, *** *P* < 0.001; **** *P* < 0.0001.

|  | n | N | Control | n | N | S1 8 d | n | N | S1 + SNc 8 d | n | N | S1 8 d+ Antag | Test | p value | Significant |
| --- | --- | --- | --- | --- | --- | --- | --- | --- | --- | --- | --- | --- | --- | --- | --- |
| In vivo electrophysiology | | | | | | | | | | | | | | | |
| **Amplitude (pA)** | | |  | | | | | | | | | | | | |
| D1 | 27 | 17 | 2.45 ± 0.41 | 14 | 5 | 6.39 ± 1.26 | 15 | 7 | 6.47 ± 1.1 | 17 | 7 | 2.56 ± 0.50 | One-way ANOVA | < 0.0001 | **** |
| *control vs S1 8d* | | | | | | | | | | | | | *Sidak's post-hoc test* | 0.0027 | ** |
| *control vs S1+SNc 8d* | | | | | | | | | | | | | *Sidak's post-hoc test* | 0.0017 | ** |
| *control vs S1 8 d + Antag* | | | | | | | | | | | | | *Sidak's post-hoc test* | 0.999 | n.s. |
| *S1 8d vs S1+SNc 8d* | | | | | | | | | | | | | *Sidak's post-hoc test* | > 0.999 | n.s. |
| *S1 8d vs S1 8d + Antag* | | | | | | | | | | | | | *Sidak's post-hoc test* | 0.009 | ** |
| *S1 SNc 8d vs S1 8d + Antag* | | | | | | | | | | | | | *Sidak's post-hoc test* | 0.007 | ** |
| D2 | 29 | 17 | 1.53 ± 0.23 | 13 | 5 | 4.98 ± 0.95 | 15 | 10 | 6.40 ± 0.87 | 11 | 6 | 4.60 ± 0.75 | One-way ANOVA | < 0.0001 | **** |
| *control vs S1 8d* | | | | | | | | | | | | | *Sidak's post-hoc test* | 0.0006 | *** |
| *control vs S1+SNc 8d* | | | | | | | | | | | | | *Sidak's post-hoc test* | < 0.0001 | **** |
| *control vs S1 8 d + Antag* | | | | | | | | | | | | | *Sidak's post-hoc test* | 0.0054 | ** |
| *S1 8d vs S1+SNc 8d* | | | | | | | | | | | | | *Sidak's post-hoc test* | 0.45 | n.s. |
| *S1 8d vs S1 8d + Antag* | | | | | | | | | | | | | *Sidak's post-hoc test* | 0.98 | n.s. |
| *S1 SNc 8d vs S1 8d + Antag* | | | | | | | | | | | | | *Sidak's post-hoc test* | 0.28 | n.s. |

**Table S10. Summary table of in *vivo* recordings (Fig. 5)**

N refers to the number of mice, n refers to the number of cells. All data are presented as mean ± SEM. N.s., non-significant; ** *P* < 0.01; ****P* < 0.001; **** *P* < 0.0001.

|  |  | | | | N | Saline | | N | Antagonists | Test | p value | Significant |
| --- | --- | --- | --- | --- | --- | --- | --- | --- | --- | --- | --- | --- |
|  | Behavior | | | | | | | | | | | |
|  | Locomotion | | | | 27 |  | | 17 |  | RM Two-way ANOVA | injection p < 0.0001 | **** |
|  |  |  |  |  |  |  |  |  |  |  | time: p < 0.0001 | **** |
|  |  | | | n | N | D1 | n | N | D2 | Test | p value | Significant |
|  | In vivo electrophysiology | | | | | | | | | | | |
|  | Resting potential (mV) | | | | |  | | | | RM Two-way ANOVA | injection p = 0.07 | n.s. |
|  |  |  |  |  |  |  |  |  |  |  | cell type: p = 0.45 | n.s. |
|  | S1 8 days | | | 15 | 5 | -76.2 ± 1.2 | 19 | 8 | -78.3 ± 0.8 |  | | |
|  | S1 8 days + antagonist | | | 15 | 5 | -82.5 ± 1.2 | 15 | 7 | -77.7 ± 2 |  | | |
|  | Membrane resistance (MΩ) | | | | |  | | | | RM Two-way ANOVA | injection p = 0.21 | n.s. |
|  |  |  |  |  |  |  |  |  |  |  | cell type: p = 0.24 | n.s. |
|  | S1 8 days | | | 15 | 5 | 100.1 ± 9.5 | 19 | 8 | 110.9 ± 7.5 |  | | |
|  | S1 8 days + antagonist | | | 15 | 5 | 111.6 ± 6.8 | 15 | 7 | 119.2 ± 6.9 |  | | |
|  | Rheobase (pA) | | | | |  | | | | RM Two-way ANOVA | injection p = 0.10 | n.s. |
|  |  |  |  |  |  |  |  |  |  |  | cell type: p < 0.0001 | **** |
|  | S1 8 days | | | 15 | 5 | 553.3 ± 32.5 | 19 | 8 | 332.1 ± 29.1 | Sidak's post-hoc test | < 0.0001 | **** |
|  | S1 8 days + antagonist | | | 15 | 5 | 493.4 ± 25.4 | 15 | 7 | 301.7 ± 19.9 | Sidak's post-hoc test | < 0.0001 | **** |
|  |  | | | n | N | S1 8 days | n | N | S1 8 days + antagonist | Test | p value | Significant |
|  | PPR D1 | | | 14 | 5 |  | 17 | 8 |  | RM Two-way ANOVA | injection p = 0.0009 | *** |
|  |  |  |  |  |  |  |  |  |  |  | stim number: p = 0.34 | n.s. |
|  | *2nd* | | |  | | 0.8 ± 0.05 |  | | 1.49 ± 0.20 | *Sidak's post-hoc test* | 0.19 | n.s. |
|  | *3rd* | | |  | | 0.76 ± 0.05 |  | | 1.97 ± 0.24 | *Sidak's post-hoc test* | 0.003 | ** |
|  | *4th* | | |  | | 0.676 ± 0.05 |  | | 2.04 ± 0.38 | *Sidak's post-hoc test* | 0.0008 | *** |
|  | *5th* | | |  | | 0.669 ± 0.04 |  | | 2.00 ± 0.40 | *Sidak's post-hoc test* | 0.001 | ** |
|  | PPR D2 | | | 13 | 5 |  | 11 | 6 |  | RM Two-way ANOVA | injection p = 0.0037 | ** |
|  |  |  |  |  |  |  |  |  |  |  | stim number: p = 0.24 | n.s. |
|  | *2nd* | | |  | | 0.85 ± 0.07 |  | | 1.25 ± 0.15 | *Sidak's post-hoc test* | 0.05 | n.s. |
|  | *3rd* | | |  | | 0.84 ± 0.09 |  | | 1.21 ± 0.14 | *Sidak's post-hoc test* | 0.08 | n.s. |
|  | *4th* | | |  | | 0.76 ± 0.08 |  | | 1.14 ± 1.3 | *Sidak's post-hoc test* | 0.07 | n.s. |
|  | *5th* | | |  | | 0.65 ± 0.06 |  | | 1.18 ± 0.16 | *Sidak's post-hoc test* | 0.005 | ** |
|  | n | N | Control | n | N | S1 8 days | n | N | S1 + SNc 8 days | Test | p value | Significant |
| PPR D1 | 27 | 17 |  | 14 | 5 |  | 15 | 7 |  | RM Two-way ANOVA | condition p = 0.056 | n.s. |
|  |  |  |  |  |  |  |  |  |  |  | stim number: p = 0.003 | ** |
| *2nd* | | | 1.03 ± 0.08 |  | | 0.8 ± 0.05 |  | | 0.92 ± 0.06 |  | | |
| *3rd* | | | 1.01 ± 0.07 |  | | 0.76 ± 0.05 |  | | 0.87 ± 0.08 |  | | |
| *4th* | | | 0.92 ± 0.08 |  | | 0.676 ± 0.05 |  | | 0.82 ± 0.09 |  | | |
| *5th* | | | 0.90 ± 0.08 |  | | 0.669 ± 0.04 |  | | 0.79 ± 0.08 |  | | |
| PPR D2 | 29 | 17 |  | 13 | 5 |  | 11 | 6 |  | RM Two-way ANOVA | condition p = 0.22 | n.s. |
|  |  |  |  |  |  |  |  |  |  |  | stim number: p = 0.028 | * |
| *2nd* | | | 1.00 ± 0.11 |  | | 0.85 ± 0.07 |  | | 0.80 ± 0.04 |  | | |
| *3rd* | | | 1.07 ± 0.14 |  | | 0.84 ± 0.09 |  | | 0.79 ± 0.05 |  | | |
| *4th* | | | 0.95 ± 0.11 |  | | 0.76 ± 0.08 |  | | 0.76 ± 0.05 |  | | |
| *5th* | | | 0.89 ± 0.12 |  | | 0.65 ± 0.06 |  | | 0.73 ± 0.05 |  | | |

**Table S11. Summary table of in *vivo* recordings (Fig. S7)**

N refers to the number of mice, n refers to the number of cells. All data are presented as mean ± SEM. N.s., non-significant; ** *P* < 0.01; ****P* < 0.001; **** *P* < 0.0001.
